# Watching ion-driven kinetics of ribozyme folding and misfolding caused by energetic and topological frustration one molecule at a time

**DOI:** 10.1101/2023.02.06.527349

**Authors:** Naoto Hori, D. Thirumalai

**Affiliations:** Department of Chemistry, University of Texas, Austin, TX 78712, USA; School of Pharmacy, University of Nottingham, Nottingham, UK; Department of Physics, University of Texas, Austin, TX 78712, USA

## Abstract

Folding of ribozymes into well-defined tertiary structures usually requires divalent cations. How Mg^2+^ ions direct the folding kinetics has been a long-standing unsolved problem because experiments cannot detect the positions and dynamics of ions. To address this problem, we used molecular simulations to dissect the folding kinetics of the *Azoarcus* ribozyme by monitoring the path each molecule takes to reach the folded state. We quantitatively establish that Mg^2+^ binding to specific sites, coupled with counter-ion release of monovalent cations, stimulate the formation of secondary and tertiary structures, leading to diverse pathways that include direct rapid folding and trapping in misfolded structures. In some molecules, key tertiary structural elements form when Mg^2+^ ions bind to specific RNA sites at the earliest stages of the folding, leading to specific collapse and rapid folding. In others, the formation of non-native base pairs, whose rearrangement is needed to reach the folded state, is the rate-limiting step. Escape from energetic traps, driven by thermal fluctuations, occurs readily. In contrast, the transition to the native state from long-lived topologically trapped native-like metastable states is extremely slow. Specific collapse and formation of energetically or topologically frustrated states occur early in the assembly process.

**Graphical abstract:** 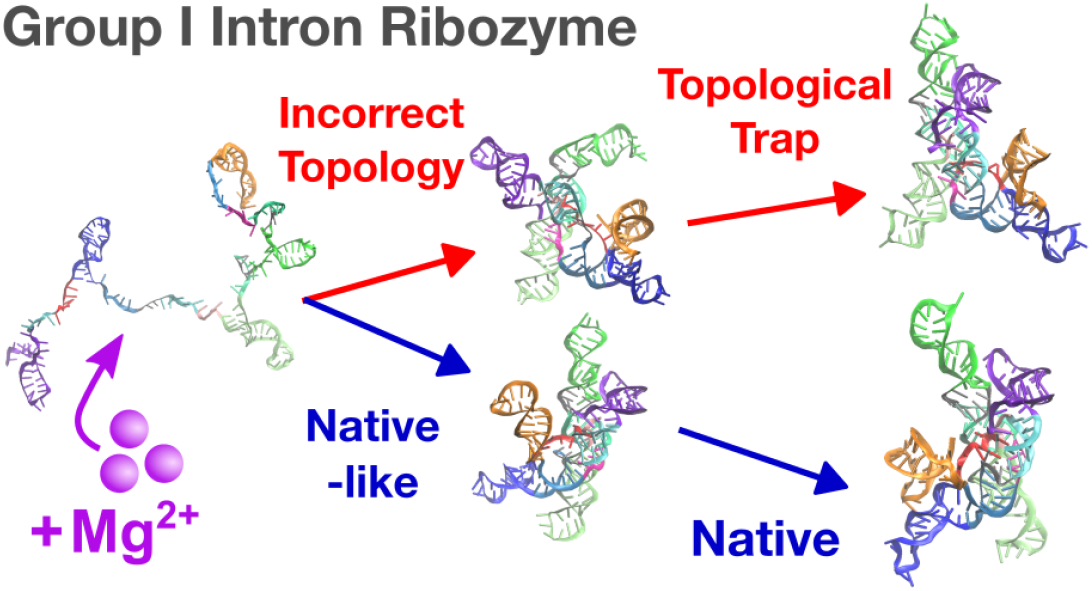

## INTRODUCTION

Many functional RNA molecules fold to specific tertiary structures, in which divalent cations play crucial roles [1–7]. Their folding pathways often consist of multiple steps covering a spectrum of time scales, and traversal through multiple pathways [8, 9]. The structural ensemble of folding intermediates is, therefore, highly heterogeneous [10]. In spite of advances in experimental methods, there are still limitations to the spatial and temporal resolution in dissecting the folding of large RNA molecules, especially the mechanisms by which divalent cations modulate the folding landscape.

Group I intron is a self-splicing ribozyme, that has been widely used in studies of RNA folding, typically either from purple bacteria *Azoarcus* or ciliates *Tetrahymena* [11]. In the presence of divalent cations (Mg^2+^), the RNA molecule folds to a specific tertiary structure (Fig. 1a-b) [12]. It is known that the folding takes place in a hierarchical manner [13, 14]. Mg^2+^ is indispensable not only for its catalytic activity but also for the formation of the native conformation [15, 16]. The double-strand helices in the core region can fold at Mg^2+^ ∼0.2 mM, whereas the complete tertiary structure requires at least ∼2 mM Mg^2+^. Woodson and coworkers revealed, by time-resolved Small Angle X-ray Scattering (tSAXS) experiments, up to 80% fraction of unfolded *Azoarcus* ribozyme reach compact structures in less than 1 ms upon the addition of 5 mM Mg^2+^ [14]. The overall folding time has been estimated to be 5 to 50 ms for the major fraction of RNA in ensemble experiments, which is faster in *Azoarcus* than *Tetrahymena* ribozyme [17], because *Azoarcus* ribozyme has smaller and simpler peripheral domains. On the other hand, it has also been known that a certain fraction of the molecule is trapped in an intermediate state, presumably because of misfolding. This persistent intermediate state needs times on the order of minutes to hours to fold to the native conformation [18].

**FIG. 1.**
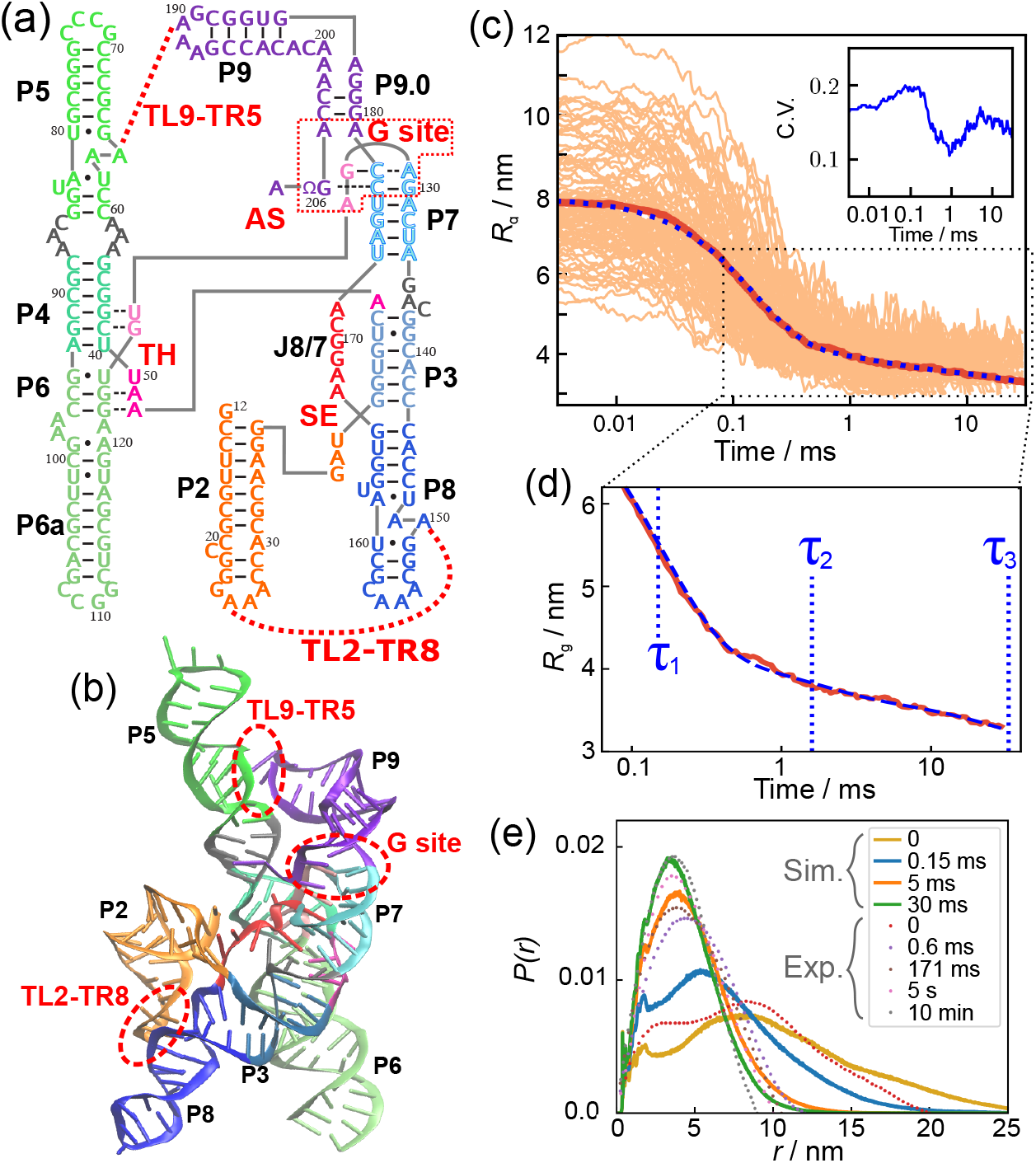
Structure and Mg^2+^-mediated collapse of the *Azoarcus* ribozyme. **(a)** The secondary structure map shows that several helices are ordered along the sequence. The helices of the *Azoarcus* ribozyme are conventionally denoted as P2 through P9 (labeled in black). P2, P5, P6, P8, and P9 are hairpin structures in which adjacent segments form the double strands, whereas P3, P4, and P7 are double strands formed by non-local pairs of segments. Several key elements involving tertiary interactions are shown in red: TH, triple helix; SE, stack exchange; G site, Guanosine-binding site; TL2-TR8, tetraloop 2 and tetraloop-receptor 8; TL9-TR5, tetraloop 9 and tetraloop-receptor 5; and AS, the active site. **(b)** Tertiary structure, taken from PDB 1U6B [12]. The same colors are used as in (a). **(c)** Time dependence of the radius of gyration (*R*_g_) averaged over 95 trajectories (thick red line). Thin lines show individual trajectories. At the beginning of the simulations *t* = 0, the average *R*_g_ is ⟨*R*_g_⟩ ≈ 7.8 nm, corresponding to the value at equilibrium in the absence of Mg^2+^. The fit using three exponential functions (see the main text) is shown by the blue dotted line. Inset: Coefficient of variation (C.V.), calculated as 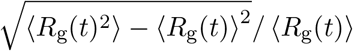 where ⟨⟩ is the average over the trajectories. **(d)** The middle to late stages of compaction are magnified from the data in (c). The three time constants are indicated by blue vertical lines. **(e)** Distance distribution functions reveal the stages in the collapse kinetics. The distribution functions were calculated using snapshots from the 95 trajectories at *t* = 0, 0.15, 5, and 30 ms. The dotted lines are experimental tSAXS data from Ref. [14].

Here, we simulate the multistep folding kinetics of *Azoarcus* group I intron by extensive Brownian dynamics simulations using a coarse-grained RNA with explicit ions [19]. In previous studies, we showed that the model reproduces Mg^2+^-concentration dependence of the *Azoarcus* ribozyme folding and correctly predicts binding sites of Mg^2+^ in equilibrium simulations [19, 20]. Here, we focused on the kinetics of the same RNA. We conducted 95 folding simulations triggered by adding 5 mM Mg^2+^ to unfolded ribozyme prepared in the absence of divalent cations. Among them, 55 trajectories reached the native conformation within the simulation time. We found that a certain fraction of simulated trajectories were trapped in misfolded states. The folding reaction took place through multiple phases as monitored by the time-dependent changes in the overall size (*R*_g_), and formation of key interactions. Most (∼80%) of secondary structures folded rapidly within the first phase. In contrast, about half of tertiary interactions formed gradually during the first and the middle phases, and the other half folded in the last phase. Non-native base pairs contributed to a manifold of metastable states comprising of a combination of mispaired helices, which slowed the folding reaction. However, one of the misfolded states mostly consisted of native interactions without mispaired helices (a topological trap). Thus, not only non-native base pairs but also the topology of the chain are relevant in characterizing the rugged RNA folding landscape. We also analyzed the dynamics of Mg^2+^ ions, and showed that Mg^2+^ rapidly replaces K^+^ when the folding reaction is initiated. Nearly 90% of the number of Mg^2+^ ions were condensed onto the RNA in the earliest phase, in which most tertiary interactions and some helices were still unfolded. Comparison of our results, with several experimental data, including time-dependent *R*_g_ from tSAXS experiments and hydroxyl radical footprinting, shows near quantitative agreement. This allows us to investigate the detailed structural changes triggered by Mg^2+^ as the *Azoarcus* ribozyme folds, events that cannot be accessed by ensemble or single molecule experiments.

## MATERIALS AND METHODS

### Three-Interaction-Site (TIS) model with Explicit ions

In order to simulate the long time scale needed to fold the ribozyme, we used the TIS model in the presence of Mg^2+^ as well as K^+^ that is in the buffer [19]. The TIS model for nucleic acids has three interaction sites for each nucleotide, corresponding to the phosphate, sugar, and base moiety (Fig. S1) [21]. All the ions are explicitly treated, whereas water is modeled implicitly using a temperature-dependent dielectric constant. The force field in which the physicochemical nature of RNA was carefully considered is given as *U*_TIS_ = *U*_bond_ +*U*_angle_ +*U*_EV_ +*U*_HB_ +*U*_ST_ + *U*_ele_. The first two terms, *U*_bond_ and *U*_angle_ ensure the connectivity of the bases to the ribose backbone with appropriate bending rigidity. The next term, *U*_EV_, accounts for excluded volume effects, which essentially prevent overlap between the beads. Hydrogen-bonding and stacking interactions are given by *U*_HB_ and *U*_ST_, respectively. We consider hydrogen bonds for all possible canonical pairs of bases (any G-C, A-U, or G-U base pairs can be formed), as well as tertiary hydrogen bonds that are formed in the crystal structure (PDB 1U6B). The stacking interactions are applied to any two bases from consecutive nucleotides along the sequence, as well as tertiary base stacking in the crystal structure. Parameters in these terms are optimized so that the model reproduces the thermodynamics of nucleotide dimers, several types of hairpins, and pseudoknots [22]. It should be emphasized that *no parameter* in the energy function was adjusted to achieve agreement with experiments on the simulated ribozyme. Thus, the results are emergent consequences of direct simulations of the *transferable TIS model*. The detailed functional forms for these terms are given in the Supplementary Methods and Table S1 in the Supplementary Data. The optimized parameters and a list of tertiary interactions can be found in the main text and supplemental information given elsewhere [19]. We showed previously that the model reproduced the experimental thermodynamics data for several RNA motifs, such as hairpin and pseudoknot, thermodynamics of *Azoarcus* ribozyme as a function of Mg^2+^ concentrations, and more recently, the thermodynamics of assembly of the central domain of the ribosomal RNA [23]. A crystal structure of *Azoarcus* group I intron (PDB 1U6B [12], Fig. 1b) was used as the reference structure for the native conformation. The nucleotides are numbered from 12 through 207 following the convention in the literature of *Azoarcus* group I intron.

### Simulation protocol

To generate the unfolded state ensemble, we performed equilibrium simulations in the absence of Mg^2+^ with 12 mM KCl, corresponding to the concentration in the Tris buffer. To enhance the efficiency of sampling the configurations of the system, we employed under-damped Langevin dynamics [24] simulations by setting the friction coefficient to 1% of water viscosity. We recorded the conformations of RNA and ions every 10^8^ steps, to be used as initial coordinates for the folding simulations. The duration of the equilibrium simulation is sufficiently long that all the initial structures are well separated in the configurational space. To generate 95 initial configurations, we ran over 3 × 10^11^ steps constituting equilibrium simulations. Brownian dynamics simulations [25] were performed to trace the folding reactions starting from an initial state in which the ribozyme is devoid of tertiary structures. The viscosity was set to the value of water, 8.9 × 10^−4^ Pa·s. The simulations were performed in a cubic 35 nm box containing ions and RNA. To minimize finite-size effects, we used periodic boundary conditions. The temperature was set to *T* = 37°C. Additional details are described in Supplementary Methods.

Starting from the initial conformations, prepared at [Mg^2+^] = 0, we triggered the folding reaction by adding 5 mM Mg^2+^. Both experiments and previous simulations [19] have shown that a solution containing 5 mM Mg^2+^ and 12 mM K^+^ drives *Azoarcus* ribozyme to structures that are catalytically active [14]. We generated 95 folding trajectories until the ribozyme reaches the folded state, or the simulation time is ≈ 30 ms. We assessed whether the ribozyme is folded to the correct native state by calculating the root-mean-square-deviation (RMSD) from the crystal structure. If the RMSD to the native structure is less than 0.6 nm, then the ribozyme is folded. We confirmed that if RMSD < 0.6 nm, all the secondary and tertiary interactions are correctly formed. Note that the experimentally (tSAXS) accessible quantity, the radius of gyration (*R*_g_), alone is not an accurate order parameter to distinguish the native structure from the misfolded structures because some of the non-native structures have *R*_g_ values that are close to the native structure. We use several measures, as described here and in the Supplementary Methods, to monitor the order of events during the folding process.

### Clustering analysis

To classify the compact conformations of the ribozyme and examine if there are any non-native (misfolded) conformations, we performed a clustering analysis. First, from all conformations in the 95 folding trajectories, we collected compact structures whose *R*_g_ ≤ 3.5 nm, regardless of their similarities to the native conformation. This criterion generated 2,162,299 structures. After reducing the number of structures to 10,811 (1/200) by random selection, a clustering analysis was done by Ward’s method using the Distribution of Reciprocal of Interatomic Distances (DRID) as the similarity measure [26].

### Ion condensation and binding

To make a quantitative comparison of condensation of monovalent (K^+^) and divalent (Mg^2+^) cations, we counted the number of ions condensed onto the RNA at each time frame. We consider that an ion is condensed when it is in the vicinity of any phosphate site in the RNA. To compare Mg^2+^ and K^+^ on equal footing, we used the Bjerrum length (*l*_*B*_ = 0.73 nm) as the cutoff distance for ion condensation. At the distance *l*_*B*_ the thermal energy balances the Coulomb attraction between a cation and an anion.

To detect tightly-bound Mg^2+^ ions, in a manner consistent with the previous study [19], we computed the contact Mg^2+^ concentration, *c*^∗^, as follows. For every phosphate site in each snapshot of the simulations, we counted the number of Mg^2+^ located in the range, *r*_0_ − Δ*r < r < r*_0_ + Δ*r*, where *r* is the distance from the phosphate, *r*_0_ = *R*_P_ + *R*_Mg_ = 0.44 nm is the sum of the excluded-volume radii of the phosphate and Mg^2+^ ion, and Δ*r* = 0.15 nm is a tolerance margin for contact (Fig. S2a). The contact Mg^2+^ concentration was then calculated by dividing the number by the spherical shell volume (Fig. S2). The definition of Mg^2+^ binding rate using the same criterion can be found in the Supplementary Methods.

### Footprinting analysis

It is known that experimental footprinting using hydroxyl radical is highly correlated with the Solvent Accessible Surface Area (SASA) of the sugar backbone [27–29]. To compare our simulation results with experimental footprinting data, we calculated SASA using FreeSASA version 2.0 [30]. Considering that hydroxyl radicals preferably cleave C4’ and C5’ atoms of RNA backbone [28], we took the larger value of the SASA of C4’ and C5’ atoms for each nucleotide. From the SASA data, we computed the “protection factor” [31] of the *i*^th^ nucleotide (see the Supplementary Methods for the detail), 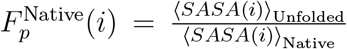 and 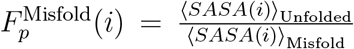 for the native and misfolded states, respectively, where the bracket indicates the ensemble average. We used the initial conformations prepared in the absence of Mg^2+^ to compute the average SASA of the unfolded state, ⟨SASA⟩_Unfolded_. In order to perform the SASA calculation, we reconstructed atomically detailed structures of the ribozyme from coarse-grained coordinates using an in-house tool (Fig. S1), which employs a fragment-assembly approach [32, 33], followed by minimization by Sander in Amber16 [34]. In Fig. S3, we show some examples of atomically detailed structures that were obtained from the conformations generated using the TIS model.

## RESULTS

There are two parts to the Mg^2+^-induced folding of the ribozyme. One pertains to the time-dependent changes in the conformations of the RNA as it folds. The second is related to the role that Mg^2+^ plays in driving the ribozyme to the folded state. It is easier to infer the major conformational changes of the ribozyme using experiments. In contrast, it is currently almost impossible to monitor the fate of the time-dependent changes in many Mg^2+^ ions as they interact with specific sites on the RNA. Both the time-dependent changes in the RNA structures and, more importantly, how correlated motions involving multiple divalent cations lead to folding, cannot be simultaneously be measured in experiments. Simulations, provided they are reasonably accurate, are best suited to probe the finer details of RNA conformational changes, and the mechanism by which Mg^2+^ drives the ribozyme to fold as a function of time. After demonstrating that our simulations nearly quantitatively reproduce the measured time-dependent changes in the size of *Azoarcus* ribozyme, we focus on the effect of Mg^2+^ on the folding reaction.

### Unfolded state ensemble is heterogeneous

We prepared the unfolded state structural ensemble in the absence of Mg^2+^ at 12 mM KCl concentration, which is the concentration in the Tris buffer [35]. The average radius of gyration, in the absence of Mg^2+^, is ⟨*R*_g_⟩ = 7.8 nm (Fig. 1c), which is in excellent agreement with experiments (≈ 7.5 nm measured by SAXS experiments in the limit of low [Mg^2+^]) [35]. Although the tertiary interactions are fully disrupted, several secondary structures, helix domains P2, P4, P5, and P8, are almost intact (Fig. S4). Nevertheless, globally the *Azoarcus* ribozyme is unstructured, with most of the characteristics of the native structure being absent (see the unfolded structures in Fig. 2). The unfolded conformational ensemble is highly heterogeneous, containing a mixture of some secondary structural elements and flexible single-stranded regions (Fig. S5).

**FIG. 2.**
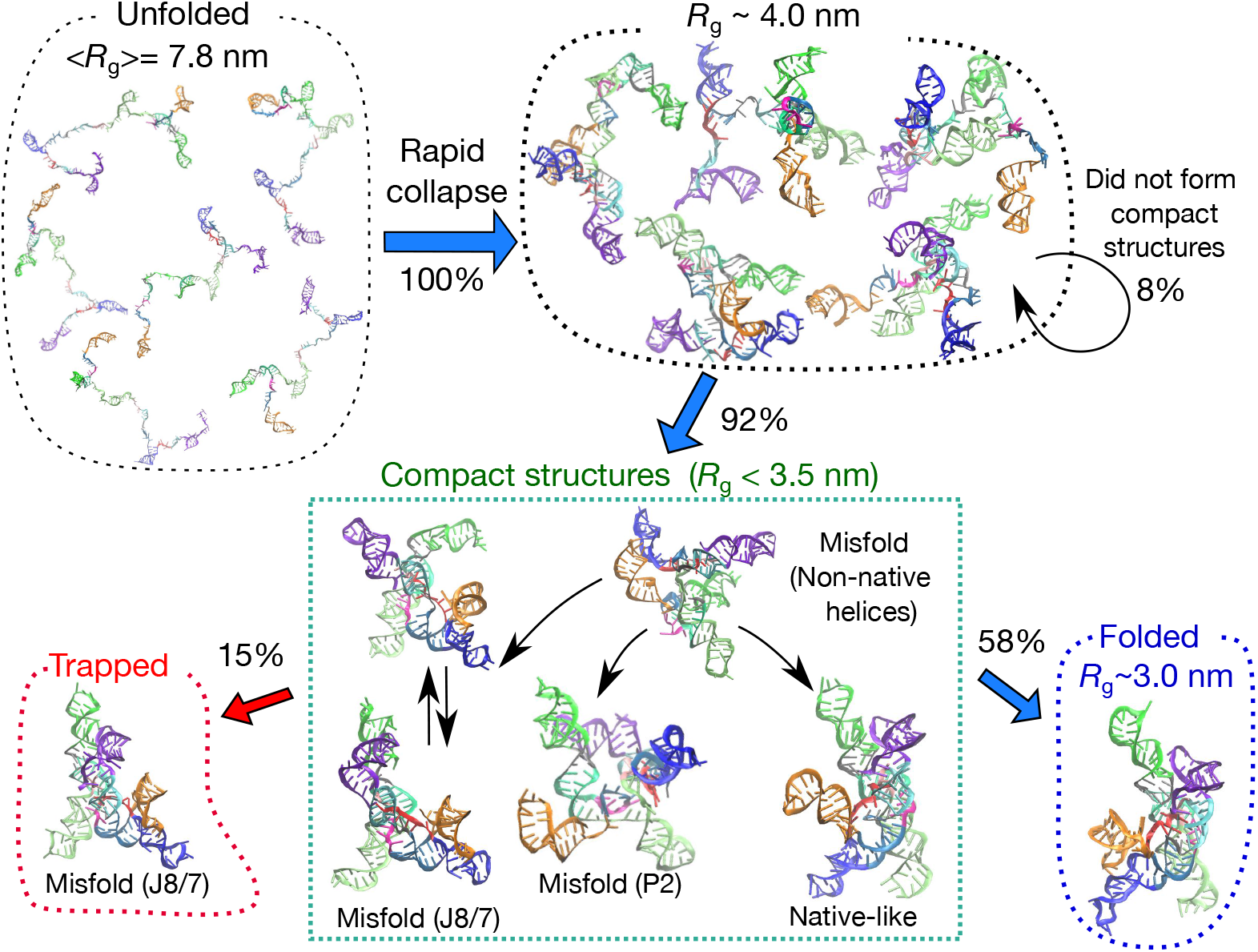
Kinetic partitioning of the trajectories. The blue and red arrows show the fate of the 95 folding trajectories initiated from the unfolded state (top left). The numbers beside the arrows indicate the fraction of trajectories in different pathways. In the compact structural ensemble, several representative misfolded structures, obtained from a clustering analysis, are shown. The structures labeled ‘Misfold (P2)’ (shown in the middle of the panel in the box) and ‘Misfold (J8/7)’ (lower left) are topological traps. Within the simulation time, 58% of the trajectories reached the folded (native) state (right bottom), whereas 15% were topologically trapped in the J8/7 misfolded state (left bottom). The remaining trajectories (27%) are trapped in the compact-structure ensemble, partly because helices are mispaired (energetic trap).

### Ribozyme collapse occurs in three stages

We first report how the folding reaction proceeds from the ensemble perspective by averaging over all the folding trajectories in order to compare with the tSAXS experiments [14]. Following the experimental protocol, we monitored the folding kinetics using the time-dependent changes in the radius of gyration, *R*_g_, describing the overall compaction of the ribozyme (Fig. 1c). At each time step, *R*_g_ was averaged over all the trajectories. At *t* → 0 (see the limit of small *t* in Fig. 1c), the average is ⟨*R*_g_⟩ ≈ 7.8 nm, corresponding to the unfolded state. The time-dependent changes in the average ⟨*R*_g_⟩ ≡ *R*_*g*_(*t*) are fit using,

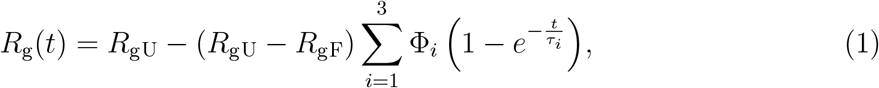

where *R*_gU_ and *R*_gF_ are the average ⟨*R*_g_⟩ of the unfolded and folded state, respectively, and *τ*_*i*_ and Φ_*i*_ are the time constants and the amplitudes (Φ_1_ + Φ_2_ + Φ_3_ = 1) associated with the *i*^th^ phase, respectively. The data could not be fit accurately using a sum of two exponential functions (Fig. S6). The best fit parameters are Φ_1_ = 0.76, Φ_2_ = 0.11, Φ_3_ = 0.13, with the corresponding time constants, *τ*_1_ = 0.15, *τ*_2_ = 1.6, *τ*_3_ = 33 ms (Fig. 1d). The three time scales describe the multi-step folding events if the radius of gyration is a reasonable order parameter for the folding reaction: (i) rapid collapse from the unfolded state (⟨*R*_g_⟩ ≈ 7.8 nm) to an intermediate state in which the RNA is compact with ⟨*R*_g_⟩ ≈ 4 nm (*τ*_*c*_ = *τ*_1_ = 0.15 ms); (ii) Further compaction to an intermediate state, *I*_*c*_, which has ⟨*R*_g_⟩ ≈ 3.5 nm; and (iii) finally folding to the native structure with ⟨*R*_g_⟩ = 3 nm. Clearly, the maximum extent of compaction occurs in the earliest stage of the folding reaction.

It is important to compare the simulation results with experimental SAXS data [14] in order to validate our model. The tSAXS results also show the three stages, with an initial decrease to ⟨*R*_g_⟩ ≈ 4 nm occurring in 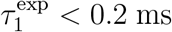. This is close to the time scale observed in the simulations, *τ*_1_ = 0.15 ms. In the tSAXS experiment, further compaction to the I_c_ state (⟨*R*_g_⟩ ≈ 3.5 nm) occurs in 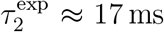, which is an order of magnitude larger than predicted in the simulations, *τ*_2_ = 1.6 ms. In the experiments [14], there is no data for *R*_g_ for time less than 0.6 ms, which might influence the estimate of 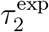. The ⟨*R*_g_⟩ of the I_c_ state is similar in the simulations and experiments (Fig. 1d).

By fitting *R*_g_ in the simulations to the three-stage kinetics, we obtained the time constant of the final transition leading to ⟨*R*_g_⟩ ≈ 3 nm, *τ*_3_ = 33 ms, that is orders of magnitude smaller than the experimental estimate 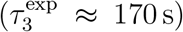. The most likely reason for this discrepancy is that, for computational reasons, we terminated the folding trajectories at 30 ms regardless of whether the RNA is folded or not. Therefore, the estimated value of *τ*_3_ from simulations is a lower bound to the time constant reported in experiments. It is worth noting that analysis of the experimental data emphasized only 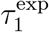 and 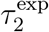, spanning times on the order of 200 ms (see Fig. 4A in [14]).

The trajectories that do not reach the native state in 30 ms undergo non-specific collapse into structures that require a longer time to reach the folded state. Nevertheless, it is clear that the simulations capture the multistage collapse of the ribozyme observed in tSAXS experiments, with near quantitative agreement for the first two phases (Fig. 1d). We confirmed that distance distribution functions at several different stages in the folding are also consistent with experiments [14] (Fig. 1e).

The amplitudes of all the three stages in the decay of *R*_g_(*t*) in the simulations are in a near quantitative agreement with fits to the measured *R*_g_(*t*) (see Figure 4 in [14]). At [Mg^2+^] = 5 mM, the calculated values from the simulations are Φ_1_ = 0.76, Φ_2_ = 0.11, and Φ_3_ = 0.13, whereas the experimental estimates are 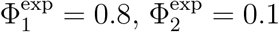, and 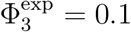. The level of agreement is remarkable, considering that no parameter in the model was adjusted to obtain agreement with any observable in the experiment. If *R*_*g*_ is a good reaction coordinate, then this would imply that nearly 80% of *Azoarcus* ribozyme folds rapidly.

The ensemble average ⟨*R*_g_⟩ hides the high degree of heterogeneity in the Mg^2+^-induced compaction of the RNA. The results in the inset of Fig. 1c show that the dispersion in *R*_g_ as a function of *t* is considerable even in the late stages of folding. This is the first indication of the importance of pathway heterogeneity in the folding process, which we further substantiate below.

### Dynamics of folding and misfolding

In Fig. 2, the fate of the trajectories and schematic folding routes are presented along with some representative structures. After the initial compaction, most trajectories (87 out of 95 ≈ 92%) show a further reduction in *R*_g_ in the second phase, in which compact structures form (*R*_*g*_ *<* 3.5 nm). Among them, 55 trajectories reach the folded state in 30 ms with structural features that are the same as in the crystal structure. To identify the distinct conformations that appear in the compact structural ensemble, we performed a clustering analysis (Supplementary Methods and Fig. S7). Although there are several misfolded elements in the ensemble structures, along with native-like structures, we identified two types of misfolding mechanisms, namely energetic traps and topological traps. The energetic trap consists of misfolded states that contain non-native helices (Fig. S8). These non-native helices can be eventually disrupted (two strands dissociate by stochastic thermal fluctuations), and undergo transition either to the native-like state or to one of the second type of misfolded states.

The second type of misfolded conformations, which we classify as topological traps, are due to frustration arising from chain connectivity [36]. In the topological trap, most of the helices are correctly formed, as in the native structure. However, the spatial arrangement of helices in some regions differs from the native structure (e.g., a part of the chain passes either on one side or the other of another strand). The incorrect topology is stabilized by several tertiary interactions, especially those involving peripheral motifs such as TL2-TR8 and TL9-TR5 (Fig. 1). As a consequence, once RNA is topologically trapped, it cannot easily escape and fold to the correct native state at any reasonable time.

We identified two distinct topological traps. In the major topological trap, ‘Misfold (J8/7)’, the J8/7 junction passes through an incorrect relative location with respect to the strands of the P3 helix (Fig. 7b). We describe this major topological trap in detail in the following sections. Although ‘Misfold (J8/7)’ resembles the native structure, the lifetime of this state is so long that it cannot be resolved even on the experimental time scale. In the minor topological trap labeled as ‘Misfold (P2)’, a part of the chain leading to the P2 helix is incorrect. This structure is stabilized by TL9-TR5 peripheral tertiary contact (See Supplementary Discussions and Figs. S9–S11).

In summary, within the simulation time of 30 ms, 55 out of 95 trajectories folded correctly to the native structure. In 14 trajectories, the ribozyme was trapped in the misfolded (J8/7) state, which is a native-like topological trap. In 18 out of the remaining 26 trajectories, compact structures formed (*R*_g_ *<* 3.5 nm) rapidly but did not fold further, partly due to the energetic or topological frustration. Because folding is a stochastic process, the times at which each trajectory (or molecule) reaches the native state vary greatly.

### Visualizing folding events in a single folding trajectory

In Fig. 3, we show a representative trajectory in which folding is completed (see also Supplementary Movie 1). In this particular case, the ribozyme reached the native structure rapidly in *t* ∼ 2.4 ms. Helices P2, P4, P5, P8, and P9 were already formed at *t* = 0, and remained intact during the folding process. From *t* = 0 to ∼ 0.5 ms, global collapse occurred, with a rapid decrease in *R*_g_ to ∼4 nm. Along with the global collapse, P7 formed at an early stage, *t* ∼ 0.15 ms. Helix P6 and the G-site tertiary interaction formed around *t* ∼ 0.4 ms. Other key interactions formed in the order of Triple Helix (TH), TL9-TR5, and P3. Note that the order of formations of these key interactions depends on the trajectory, and is by no means unique. After the formation of the P3 helix, it took a relatively long time (∼ 1.5 ms) for P2 (orange domain in Fig. 3) to find the counterpart P8 (blue). At *t* ∼ 2.4 ms, the formation of tertiary interactions between the two domains (TL2-TR8) results in the native conformation. Figs. S12 and S13 show two other examples in which the folding was completed but on a longer time scale. In the trajectory in Fig. S12, a mispaired helix in P6 formed early (*t <* 1 ms), preventing it from folding to the native state smoothly. Eventually (*t* ∼ 25 ms), the misfolded P6 was resolved, leading to the correct native fold.

**FIG. 3.**
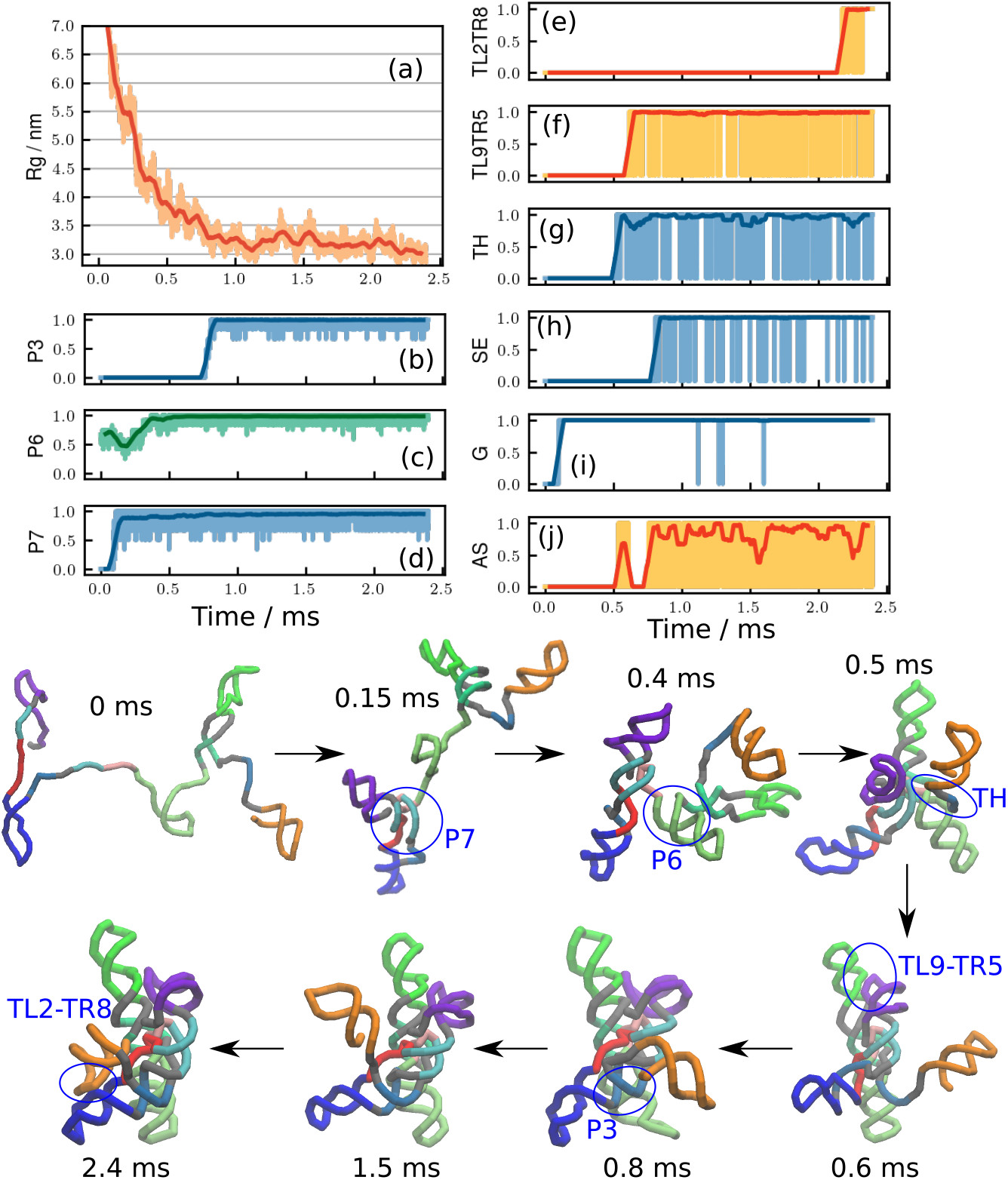
A representative rapid folding trajectory. The ribozyme folds in *t* ∼ 2.4 ms without being kinetically trapped. **(a)** Time dependent changes in *R*_*g*_. **(b-d)** Fraction of helix formations for (b) P3, (c) P6, and (d)P7. **(e-j)** Fraction of major tertiary interactions (e) TL2-TR8, (f) TL9-TR5, (g) Triple Helix, (h) Stack Exchange, (i) G site, and (j) Active Site. See Fig. 1(a-b) for the locations of these structural elements. Thin lines, with light colors are raw data, and thick lines with dark colors are averaged over 50 *μ*s window. **(Bottom)** Eight representative structures at several different time points. Major conformational changes are indicated by blue circles with labels. See Supplementary Movie 1 to watch the trajectory.

Fig. 4(a) shows a series of snapshots from a trajectory where the ribozyme is topologically trapped in the Misfold (J8/7) state. In this trajectory, two key interactions in the peripheral regions, TL2-TR8 and TL9-TR5, formed early (*t <* 1 ms). However, the junction J8/7 was in the wrong position with respect to the P3 helix strands, causing a misfolding (Fig. 7b). Because the incorrect chain topology cannot be resolved unless both of the peripheral interactions unfold, the RNA stays in this misfolded topological trap for an arbitrarily long time.

From all the folding and misfolding trajectories, we calculated the distributions of first passage times to either the folded or the trapped state. The time-dependent fractions in these two states, shown in Fig. 4(b), reveal that trajectories that reach native-like structures (*i*.*e*. either folded or topologically trapped states) by ∼5 ms fold correctly. In contrast, trajectories that take a longer time to be native-like (*>* 5 ms) are kinetically trapped. The former type of trajectories would be the consequence of the specific collapse in the earliest stage of the folding.

**FIG. 4.**
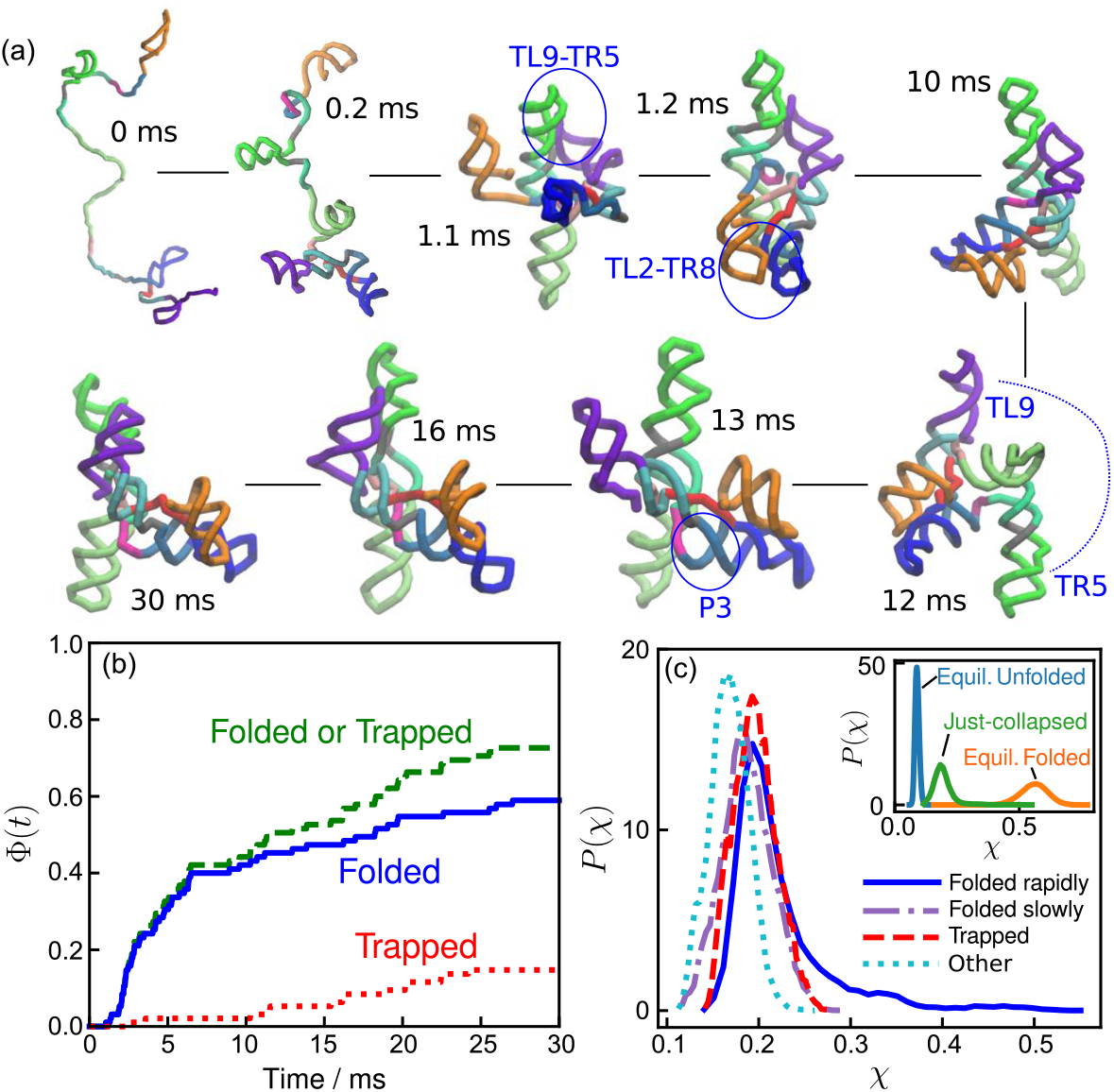
Propensity to fold correctly is determined early. **(a)** A representative misfolding trajectory showing the formation of long-lived topologically-trapped states. Two key interactions in the peripheral regions, TL2-TR8 and TL9-TR5, formed early in the folding process (*t <* 1 ms), resulting in the junction J8/7 with an incorrect topology (See Fig. 7). Because the incorrect chain topology cannot be resolved unless both of the peripheral interactions unfold, the RNA stays topologically trapped for the rest of the simulation time. See Fig. S14 for trajectories of *R*_g_, fractions of secondary and tertiary elements, Fig. S15 for another example of a kinetic trap, and Supplementary Movie 2 for watching the trajectory. **(b)** Specific collapse leads to rapid folding. From the distributions of the first passage times to the folded state,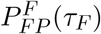, we calculated,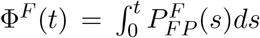 Smilarly,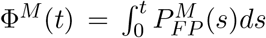,where 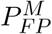 is the distribution of mean first passage times to the trapped state. **(c)** Fate of the ribozyme immediately following the initial collapse. The probability distributions of the structure overlap (*χ*) with respect to the native structure; *χ* = 0 indicates no similarity to the crystal structure, and *χ* = 1 corresponds to the native state. (Inset) The distribution of *χ* immediately after collapse (*t <* 150 *μ*s, green line, “Just-collapsed”) compared with distributions of the equilibrium unfolded state (blue, [Mg^2+^] = 0 mM) and folded state (orange, [Mg^2+^] = 5 mM). The distribution of the “just-collapsed” ensemble in the main figure is decomposed into four distributions depending on the fate of each trajectory: Folded rapidly, trajectories reached the correct folded state within 5 ms; Folded slowly, trajectories reached the correct folded state after 5 ms but within the maximum simulation time (30 ms); Trapped, trajectories where the RNA was trapped in the major misfolded state; trajectories labeled Other were neither folded nor misfolded.

### Kinetics of secondary- and tertiary-structure formation

The folded RNA structures are composed of secondary structural motifs, which are often independently stable and are consolidated by tertiary contacts to render the ribozyme compact. The most abundant elementary structural unit is the double-stranded helix (Fig. 1a). In Fig. 5, time-dependent formations of secondary and tertiary interactions, represented by average energies stabilizing these motifs, are plotted. Each interaction type is further categorized into two main chemical components, hydrogen bonding (H-bond) and base stacking. Fig. 5 (a-b) shows that most secondary structures are rapidly formed in the first phase (*t* ≲ 0.15 ms), although certain secondary interactions form only in the late stages. In contrast, the formation of most tertiary interactions occurs in the middle (*t* ∼ 1.5 ms) and the last phase (*t* ≳ 10 ms) (Fig. 5c-d). These findings illustrate the hierarchical nature of RNA folding kinetics in the ensemble picture, where formations of secondary structures are followed by tertiary contacts [37, 38].

**FIG. 5.**
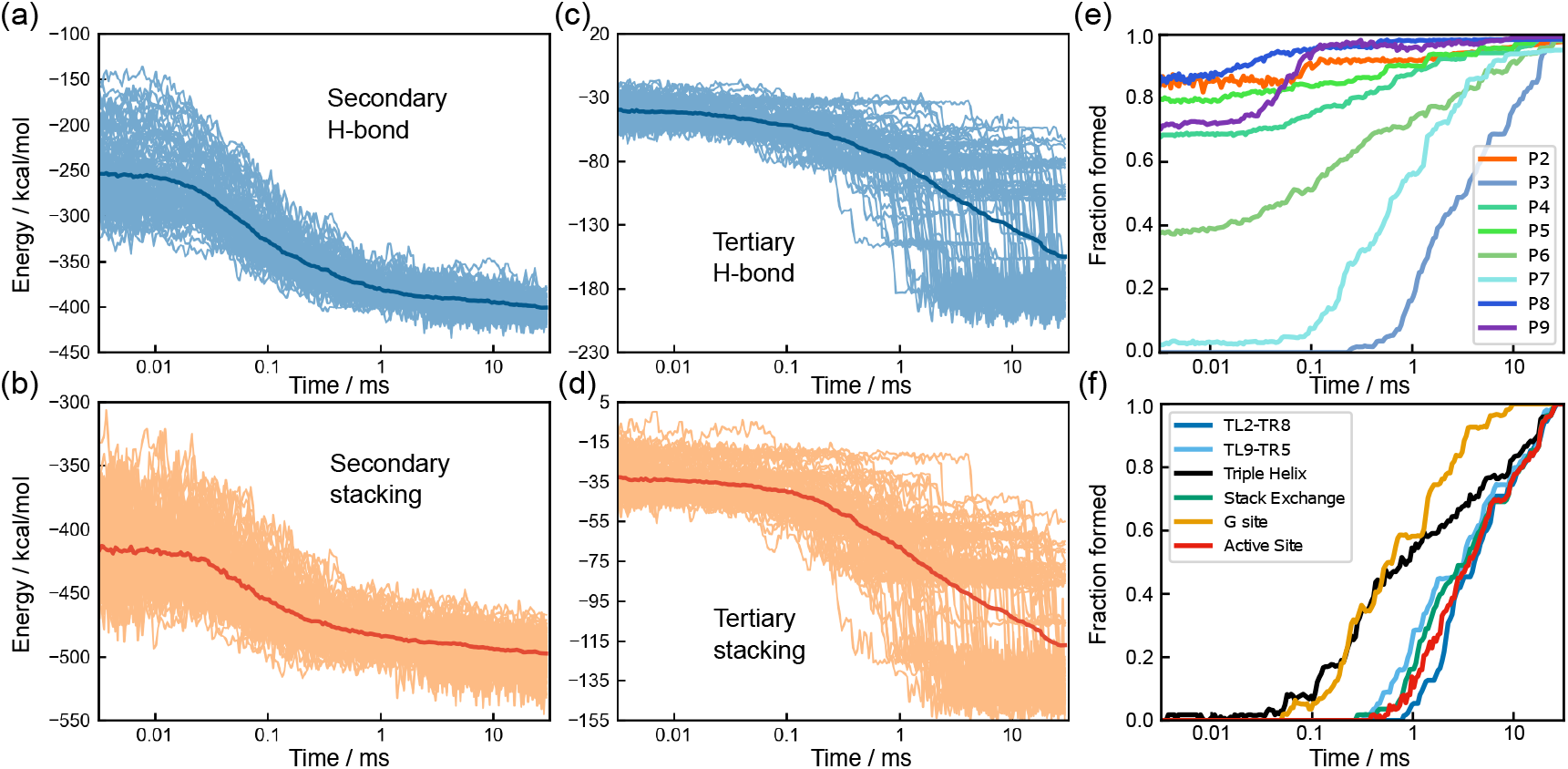
Hierarchical formation of secondary and tertiary structures. (a-d) Time-dependent formation of structural elements represented by the potential energies associated with (a) secondary hydrogen bond (H-bond), (b) secondary stacking, (c) tertiary H-bond, and (d) tertiary stacking. The thin lines are the results for the 95 individual trajectories, and the thick line in each panel is the average over all trajectories. (e) Time dependence of helix formation for helices P2 to P9. (f) Formation of six key elements associated with tertiary interactions (see Fig. 1).

We then investigated the kinetics of individual helix formations by calculating time-dependent fractions of helix formations in the folding trajectories (Fig. 5e). In summary, all helices except P3 and P7 fold early, typically in *t <* 0.1 ms. Kinetics of involving helices P3 and P7 are particularly slow because the two strands of P3 and P7 are far apart in the sequence, and thus it takes substantial time to search each other. Supplementary Discussion contains further details on the folding of individual secondary structures.

Equilibrium ensemble simulations [19] identified several key tertiary interactions (shown in red symbols in Fig. 1) by varying the Mg^2+^ concentration. In Fig. 5(f), formations of those tertiary interactions are shown as time averages over the folded trajectories. Here, we find that the triple helix (TH) and Guanosine binding site (G site) form earlier than the other key elements. This is consistent with the results of equilibrium simulations [19] that reported that TH and G sites are formed at lower Mg^2+^ concentrations compared to other key interactions. The time range of the formation corresponds to the second phase in Fig. 1(c). Following the TH and G site formation, other key interactions form, mainly in the last phase (*t >* 10 ms).

Interestingly, the kinetics of G site formation resembles the formation kinetics of the P7 helix (Fig. 5e). From the secondary structure (Fig. 1), this can be explained by noting that the G site is formed with a part of the P7 helix. Since the two strands of P7 are separated along the sequence, the formation of the P7 helix is a rate-limiting step for the formation of the G site. This is reminiscent of the diffusion-collision model proposed for protein folding [39].

### Counterion release kinetics

We now turn to the role of the cations, K^+^ and Mg^2+^, in driving the assembly of *Azoarcus* ribozyme. The interplay between the unbinding of K^+^ ions and the association of Mg^2+^ to the ribozyme as it folds is vividly illustrated in Fig. 6(a). The figure shows the time dependence of the number of cations condensed onto RNA, averaged over all the trajectories (see Methods for the definition). At *t* = 0, on average, 40 K^+^ are condensed onto the ribozyme (see Fig. S16 for snapshots). The monovalent K^+^ cations are rapidly replaced by Mg^2+^ in *t* ≲ 0.02 ms, which shows dramatically the counterion release mechanism anticipated by the application [1, 20] of the Oosawa-Manning theory [40]. In this time scale, nearly 90% of Mg^2+^ ions in the final native state are already condensed, even though most of the tertiary interactions and some helices are still unfolded. The replacement of K^+^ ions by Mg^2+^ shows that even the initiation of folding requires the reduction in the effective charge on the phosphate groups, which is accomplished efficiently by divalent cations. However, in this rapid process, some of the structures that form are topologically or energetically frustrated, thus greatly increasing the folding time. The premature condensation of Mg^2+^ has been found to produce kinetically heterogeneous structures that rearrange slowly both in Holliday junctions [41], and group II introns [42]. A fraction of unfolded RNAs undergoes specific collapse, adopting compact structures that reach the native-like fold rapidly.

**FIG. 6.**
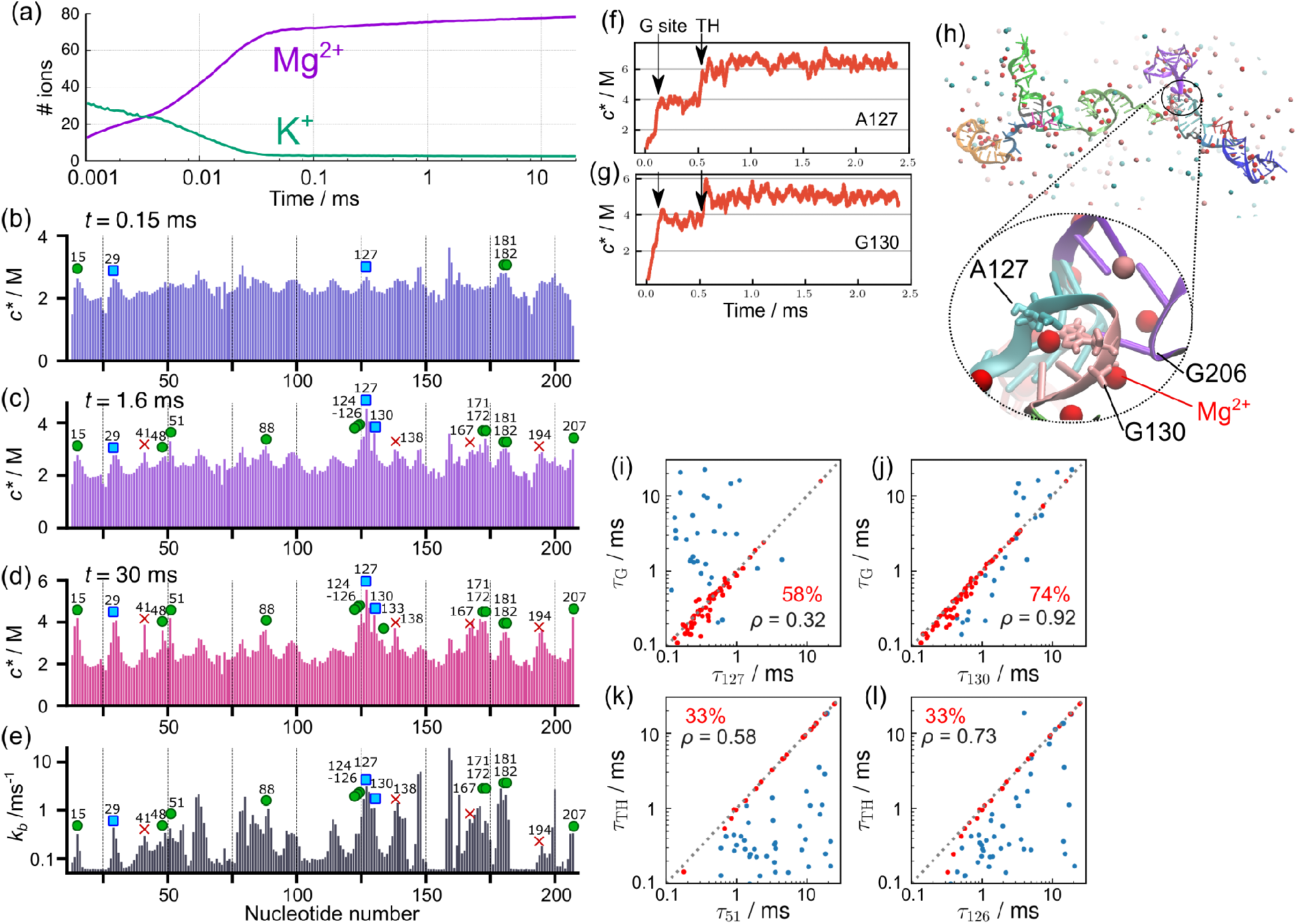
Fingerprints of Mg^2+^ association and correlation with tertiary contact formation. **(a)** Number of cations condensed onto the RNA averaged over all the trajectories. At *t* = 0, on average, there are 40 K^+^ ions in the neighborhood of the ribozyme. K^+^ cations are rapidly replaced by Mg^2+^ in *t* ≲ 0.02 ms. **(b-d)** Mg^2+^ fingerprints, measured using local nucleotide-specific concentration at (b) *t* = 0.15 ms, (c) 1.6 ms, and (d) 30 ms. Symbols indicate nucleotides that are either in direct contact with Mg^2+^ (green circles) or linked via a water molecule (cyan squares) in the crystal structure [12]. The red crosses are nucleotides predicted to bind Mg^2+^ in the equilibrium simulations [19]. **(e)** Binding rates calculated from the mean first passage times of Mg^2+^ binding event at each nucleotide. **(f, g)** Trajectories displaying contact Mg^2+^ concentration at nucleotides (f) A127 and (g) G130 taken from the folding trajectory shown in Fig. 3. The folding times of the G site and Triple Helix (TH) are indicated by black arrows. **(h)** A snapshot, at *t* = 0.11 ms, when the G site is formed in the trajectory in (f) and (g). Nucleotides A127 and G130 are shown in stick, and Mg^2+^ ions are spheres in red. **(i, j)** Scatter plots of the first passage times of Mg^2+^ binding to (i) A127 and (j) G130 versus the first passage time of the G site formation. If the Mg^2+^ binding and the G site formation occurred concurrently (|*τ*_G_ − *τ*_*i*_| *<* 0.2 ms), the two events are regarded as strongly correlated and plotted in red; otherwise it is shown in blue. In each panel, the fraction of strong correlations (red points) is indicated by the percentage in red, associated with the Pearson correlation (*ρ*) calculated using all the data points. **(k, l)** Same as (i-j) except for Mg^2+^ binding to (k) U51 and (l) U126 versus the first passage time of Triple Helix (TH) formation.

### Fingerprint of Mg^2+^ associations

Among the condensed Mg^2+^ ions, some bind at specific sites as seen in the crystal structure [12]. In Fig. 6(b-d), we show the time-dependent Mg^2+^ densities, *c*^∗^, at each nucleotide site. At many of the specific binding sites, Mg^2+^ ions associate with the ribozyme in the early stage of folding. For instance, at *t* = 0.15 ms, although most tertiary interactions are not formed (Fig. 5f), there are several peaks in the Mg^2+^ densities (Fig. 6b). Interestingly, nucleotides 15, 29, 127, and 181 are the Mg^2+^ binding sites in the crystal structure. This shows that some of the specific binding sites are occupied by Mg^2+^ at the earliest stage of folding. Coordination of Mg^2+^ at these sites is required to reduce the electrostatic penalty for subsequent tertiary structure formation. In the intermediate time scale, *t* = 1.6 ms, additional nucleotides bind Mg^2+^, as shown in Fig. 6(c). At *t* = 30 ms (Fig. 6d), we confirmed that Mg^2+^ binding sites are consistent with the crystal structure [12], which is another indication that the model is accurate.

We next investigated if there is a correlation between Mg^2+^-binding kinetics and the thermodynamics of ion association. Using the same distance criterion for Mg^2+^ binding to the RNA, we computed the mean first passage time (MFPT) of Mg^2+^ coordination to each phosphate site. The binding rate was calculated as the inverse of the MFPT and is shown in Fig. 6(e). The nucleotides that have greater binding rates (*k*_*b*_) correspond to those that have higher Mg^2+^ densities in the equilibrium simulations [19], including the positions found in the crystal structure [12]. This shows that the association of the ions to high-density Mg^2+^ sites occurs rapidly in the early stage of the folding. These results show that there is a remarkable consistency between the order of accumulation of Mg^2+^ ions at specific sites of the ribozyme and the conclusions reached based on equilibrium titration involving an increase in Mg^2+^ concentrations. The coordination of Mg^2+^ at early times to nucleotides are also the ones to which Mg^2+^ ions bind at the lowest Mg^2+^ concentration [19], thus linking the thermodynamics and kinetics of ion association to the ribozyme.

### Specific Mg^2+^ binding drives formation of tertiary interactions

From the kinetic simulation trajectories, we further analyzed whether Mg^2+^ binding events directly guide the formation of tertiary interactions. One of the key tertiary elements that folds in the early stage is the G site comprising the G-binding pocket and G206 (Fig. 1). There are two peaks corresponding to A127 and G130 in the Mg^2+^ fingerprint both in kinetics (Fig. 6e) and thermodynamics [19] simulations, consistent with a Mg^2+^ ion bound between the two nucleotides in the crystal structure [12]. We found a strong correlation between the first passage times of Mg^2+^ binding at these nucleotides, and the formation of the G site. For example, in the same trajectory, as shown in Fig. 3, Mg^2+^ binding to those nucleotides are noticeable as an increase in contact Mg^2+^ concentration (*c*^∗^) at the same time as G site formation (Fig. 6 f-g).

The scatter plots in Fig. 6(i, j) show the correlation between the first passage time for Mg^2+^ binding (*τ*_127_ and *τ*_130_) and the first passage time for G-site formation (*τ*_G_) from all the trajectories. For both A127 and G130, there is a distinct temporal correlation, reflected as dense data points in the diagonal region. In the majority of the trajectories (74% for G130 and 58% for A127), the G-site formation and Mg^2+^ binding occurred simultaneously within 0.2 ms (shown as red points in Fig 6). Overall, Mg^2+^ binding to G130 exhibited a higher correlation (Pearson coefficient *ρ* = 0.92), whereas the correlation for A127 was lower (*ρ* = 0.32) because G-site formation occurred later than the Mg^2+^ binding in some trajectories (data points on the upper left triangle in Fig. 6i). Nevertheless, for both the nucleotides, the G-site formation does not precede Mg^2+^ binding, indicated by the absence of data points on the lower right triangle. We conclude that Mg^2+^ binding to these nucleotides is a necessary condition for the formation of the G site.

Strikingly, there are also noticeable increases in the local Mg^2+^ concentration at nucleotides A127 and G130 in the later stage, corresponding to the time when another tertiary element, Triple Helix (TH), forms (also indicated by arrows in Fig. 6(f-g)). Because the G site and TH are spatially close to each other, the formation of TH further stabilizes the Mg^2+^ associations at the G site, especially A127, which is located at the end of the strand that constitutes the TH. The temporal correlations between these nucleotides and both the G site and TH show that Mg^2+^ binding to some nucleotides in the core of the ribozyme contribute to more than one tertiary contact.

Fig. 6 (k, l) show that U51 and U126 also bind Mg^2+^ ions upon folding of TH, which is consistent with the observation of two Mg^2+^ ions in the crystal structure. The scatter plots show that, in some trajectories, TH forms before Mg^2+^ ions bind to U51 and U126, indicating that Mg^2+^ binding to these nucleotides are not needed for the formation of the TH. This supports the finding that TH formed in 60% of the population even at submillimolar Mg^2+^ concentration in the equilibrium simulation study [19]. We found similar correlations for other key tertiary elements, Stack Exchange, TL2-TR8, and TL9-TR5. See Supplementary Discussion and Figs. S17–S20.

### Non-native base pairs impede folding both to the native and topologically-trapped states

Because our model allows any combination of canonical (G-C and A-U) and Wobble (G-U) base pairs to form, we found a number of non-native base pairs in the folding process. Some of these base pairs formed only transiently, whereas others have relatively long lifetimes, suggesting a link between the formation of non-native base pairs and misfolded states. By counting the frequencies of each non-native base pair, we identified five frequently mispaired strand-strand combinations (Fig. S8). These strands are parts of helices, P3, P6, P7, and junction J8/7. For instance, each strand of P3 forms a double-strand helix with another strand from P7, leading to the formation of mispaired helices P3u-P7u and P3d-P7d (Fig. S8 right top). Here, we introduced the notation, *u* and *d*, to distinguish between the two strands for native helices, the upstream (5’-end) strand *u* and the one downstream *d*. Helices, P3, P6, P7 are unfolded in the absence of Mg^2+^ (Fig. S4). Given that these mispaired helices consist of at least four consecutive base pairs, except P3d-J8/7, it is reasonable that such incorrect pairings are formed in the process of the unfolded strands searching for their counterparts. In addition, we found alternative secondary structures of P6 helix, alt-P6 (Fig. S8 bottom right).

In Fig. S21(a), we show the time-dependent fractions of mispaired helices. One of such mispairing, P3d-J8/7, formed earlier than others, and the fraction is also higher. Interestingly, there is a significant fraction of such mispaired helices even in folding trajectories that reach the native state. Indeed, the time-dependent fraction for folded and misfolded trajectories resemble each other (Fig. S21 b-c). In both cases, the fractions of mispaired helices increase between 0.1 *< t <* 1 ms, and then decrease to nearly zero at *t* ∼ 30 ms. In contrast to expectation, the persistent misfolded states do not have a significant amount of mispaired helices. We conclude that the mispaired helices (energetics traps) often form regardless of whether folding is on the pathway to the native state or kinetically trapped.

### Footprinting data is consistent with the formation of misfolded states

Hydroxyl radicals are often used to study hierarchical structure formations in footprinting experiments. In hydroxyl radical footprinting (analogous to hydrogen exchange experiments using NMR for proteins), one can detect the extent to which nucleotides in RNA are protected from cleavage reactions by hydroxyl radicals. It is known that the degree of protection is highly correlated with the solvent-accessible surface area (SASA) of the backbone sugar atoms, whereas there is no direct relationship between protection and its sequence and secondary structure [27–29]. Consequently, the technique is useful for assessing the regions that are densely packed. In the context of ribozyme folding, it is often reported as “protections”, which indicate regions where tertiary interactions form to a greater extent compared to some reference state, which is typically the unfolded state before folding is initiated. For the *Azoarcus* ribozyme, several footprinting data are available in the literature [17, 31, 46–51]. Because it is difficult to characterize the molecular details of the misfolded conformations in experiments, our simulations provide the needed quantitative insights into the protection factor at the nucleotide level, thus filling in details that cannot be resolved experimentally.

We calculated the protection factors (footprints) for the native and the trapped ensembles based on SASA values of the simulated structures, and compared them with the experimental data. The two sets of footprints from the native and misfolded simulation ensemble have a similar pattern (Fig. 7d) but with differing amplitudes, which is an indication of the extent of protection. Even though the two structural ensembles have different chain topologies, there are many well-packed regions in common. This also implies that the ensemble of misfolded conformations shares many common characteristics with the native structures, as implied by the kinetic partitioning mechanism (KPM) [36, 52].

**FIG. 7.**
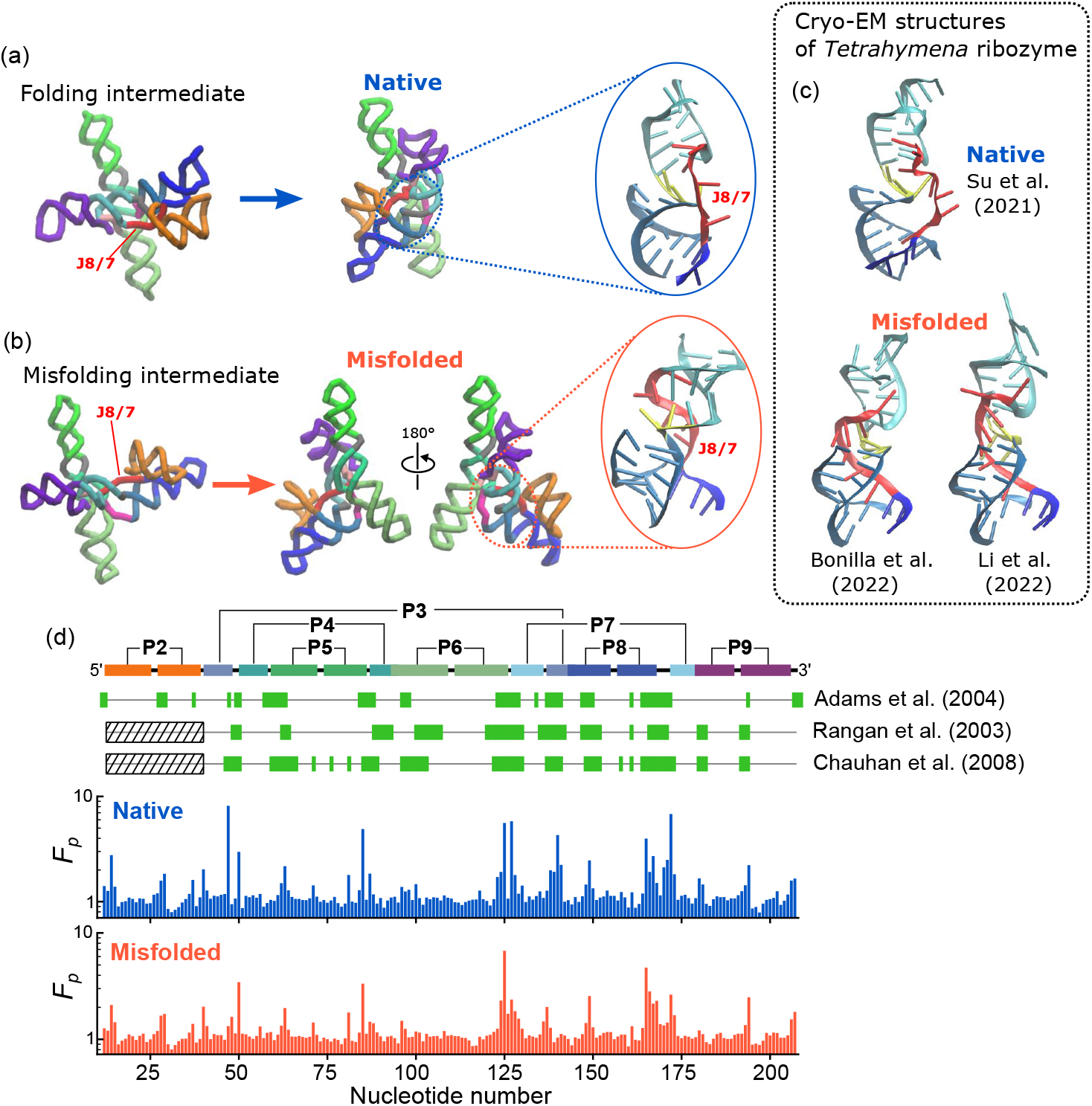
Topological frustration in the persistent metastable state. **(a-c)** The major misfolded structure (J8/7) and its intermediate (b) are compared with the native intermediate and the folded structures (a). The spatial arrangements of J8/7 and other strands at the core are depicted on the right in blue and red circles. In the misfolding intermediate (b, left), the strand J8/7 (colored in red) passes through an incorrect location relative to the other two strands of the P3 helix (cyan), resulting in a topologically trapped state. This incorrect topology cannot be resolved unless other tertiary contacts such as TL2-TR8 (contact between domains P2 (orange) and P8 (blue)) disengage, which is unlikely to occur under the folding condition. As a consequence, it leads to a metastable state that is as compact as the folded state but with an incorrect chain topology. In panel (c), the same region in the native (upper, PDB 7EZ0 [43]) and misfolded (lower, PDB 7UVT [44] and PDB 7XSK [45]) *Tetrahymena* ribozyme, solved by cryo-EM are shown for comparison. **(d)** Comparison of experimental footprinting data with SASAs calculated from simulations. Protection factor (*F*_*p*_) are calculated for the native (middle, blue) and misfolded ensembles (bottom, red) from simulations. Protected nucleotides, indicated by green-filled rectangles on top, are from data in three experiments, as labeled on the right [17, 31, 46]. The secondary structure is shown on top for reference. Note that protections of nucleotides in the P2 helix were not resolved in two experiments for technical reasons.

Experimental and calculated footprints are mostly consistent with the positions in both the protection factor profiles (Fig. 7d). There are 28 nucleotides commonly protected in the three experimental data compared [17, 31, 46]. Among these 28 nucleotides, 23 nucleotides are protected in the native ensemble (sensitivity (true positive rate) 0.82 and specificity (true negative rate) 0.79), whereas 24 nucleotides are protected in the misfolded ensemble from the simulation (sensitivity 0.86 and specificity 0.83), using a threshold protection factor, *F*_*p*_ = 1.2.

The analysis based on Fig. 7(d) quantitatively show that our simulation data and experiments are consistent with each other. However, the comparison also shows that footprinting analyses may not uniquely distinguish the native structure from the misfolded conformation unless the amplitudes are quantitatively compared. Chauhan and Woodson [17] (their footprinting data is shown in Fig. 7d) reported that about 20% of the population was mis-folded, although these experiments cannot provide the molecular details of the misfolded structures. Given the good agreement found in Fig. 7, we predict that the misfolded state identified in the simulations contributes to the 20% fraction identified in experiments. The topologically-trapped states are more flexible than the native structure, which is reflected in the decreased amplitude in the protection factor (Fig. 7d).

## DISCUSSION

We performed coarse-grained simulations to reveal the structural details and the mechanisms by which specific and correlated association of Mg^2+^ with nucleotides drive the multistep folding kinetics of *Azoarcus* group-I intron RNA. The simulated collapse kinetics (*R*_*g*_ versus time) and the tSAXS experimental data [14] are in excellent agreement with each other. There are three major phases in the folding kinetics: (1) Rapid collapse from the unfolded (⟨*R*_g_⟩ ≈ 7.8 nm) to an intermediate state in which the RNA is compact (⟨*R*_g_⟩ ≈ 4 nm). (2) The second phase involves the formation of the I_c_ state that is almost as compact as the native structure (⟨*R*_g_⟩ ≈ 3.5 nm). (3) In the final phase, there is a transition to the native structure with ⟨*R*_g_⟩ ≈ 3 nm. Interestingly, most (about 80%) of the secondary structures are formed within the first phase. In contrast, only about half of the tertiary interactions form incrementally during the first and second phases, and the remaining tertiary contacts form in the last phase. This suggests that the folding transition state is close to the folded state, which confirms the conjecture made previously [53].

A surprising finding in our simulations is that Mg^2+^ ions condense onto the ribozyme over a very short time window, *t <* 0.05 ms, which results in the release of K^+^, an entropically favorable event. Nearly 90% of Mg^2+^ ions are condensed in this time frame, even though most of the tertiary interactions and some helices are disordered. These findings show that Mg^2+^ condensation, in conjunction with K^+^ release, precedes the formation of major ion-driven rearrangements in the ribozyme. We believe that this is what transpires in ribozymes and compactly folded RNAs.

### Kinetic partitioning

The initial ribozyme collapse could be either specific, which would populate native-like structures that would reach the folded state rapidly, as predicted by the KPM [36, 54], or it could be non-specific. In the latter case, the ribozyme would be kinetically trapped in the metastable structures for arbitrarily long times. In either case, theory has shown that the collapse time, *τ*_*c*_ ≈ *τ*_0_*N*^*α*^ (*N* is the number of nucleotides and *α* ≈ 1) with the prefactor, *τ*_0_, that is on the order of (0.1 − 1)*μ*s [54]. Taking *τ*_0_ ≈ 0.5 *μ*s leads to the theoretical prediction that for the 195-nucleotide *Azoarcus* ribozyme, *τ*_*c*_ ≈ (0.02−0.2)ms, which is in accord with both simulations and experiments.

To determine if the structural variations at the earliest stage of folding affect the fate of the RNA, we analyzed the ensemble of conformations immediately after the initial collapse (*t <* 150 *μ*s). Fig. 4(c) shows the probability distributions of the structural overlap function (*χ*) calculated for the *just-collapsed* ensemble. The order parameter, *χ*, measures the extent of structural similarity to the native structure (0 for no similarity and 1 if it matched the folded state). The data is decomposed into four categories depending on the fate of each trajectory, rapidly folded, slowly folded, trapped, and trajectories that are neither folded nor kinetically trapped. There are two major findings in this plot: (1) Trajectories that result in either folded or trapped states have higher structural similarity to the folded state compared to those that do not reach these states at an early folding stage. (2) The distribution of the rapidly folded ensemble has a long tail with substantial similarity to the native structure, indicating that rapid folding to the folded structure arises from specific collapse. Although this result was expected on theoretical grounds [55], which has been established for RNA molecules whose folding rates vary over 7 orders of magnitude [56], there has been no direct demonstration of specific collapse and the associated structures until this study.

### Mispaired secondary structures impede folding

We observed several incorrect base parings en route to either the native or the misfolded state (Figs. S8 and S21). RNA sequences, in general, tend to form diverse secondary structures because there are likely multiple pairs of partially complementary regions. For instance, in mRNAs that do not have specific tertiary structures, many different patterns of secondary structures have been observed for a single sequence [57]. It is clear that even for a well-evolved sequence that has a specific tertiary structure, like the ribozyme, non-native complementary pairs can not be entirely avoided [58]. Such mispaired helices inevitably slow down the folding [59] and have to unfold, at least partially, before the RNA reaches the native state [60].

For group I intron ribozyme, it has been suggested that helix P3 has an alternative paring pattern (alt-P3) [61], which we observe as P3d-J8/7 in the simulations. The alt-P3 is thought to be a major reason for the slow folding rates of group I intron ribozyme. In accord with this proposal, it was shown that a point mutation in alt-P3, which stabilizes the correct pairing, increased the folding rate by 50 times in *Tetrahymena* ribozyme [62]. In line with the same reasoning, *Candida* ribozyme, which does not have stable alt-P3, folds rapidly without kinetic traps [63]. To our knowledge, other combinations of mispaired helices reported here (Fig. S8) have not been detected in experiments, even though such mispaired helices are permissible. There are a variety of metastable structures that render the folding landscape of RNA rugged, besides alt-P3.

### Topological frustration causes trapping in long-lived metastable states

We also found that the intermediate state, *I*_*c*_, consists of not only on-pathway conformations but also misfolded structures. The *Azoarcus* ribozyme folds correctly in a shorter time than topologically similar but larger ribozymes such as group I intron from *Tetrahymena*. Nevertheless, several experiments have shown the existence of metastable states [14, 18, 49]. These metastable states slowly transition to the native state, which can be accelerated by urea [8, 64]. We did not see refolding events from the misfolded state to the native state within our simulation time (30 ms). This is also consistent with experiments, which showed that the misfolded state is long-lived and remained stable after 5 minutes of incubation with Mg^2+^ [18]. The estimated refolding rate was 0.29 min^−1^ at 32°C with 5 mM Mg^2+^.

In our simulations, the ribozyme misfolded to a topologically frustrated state when one of the peripheral contacts, TL2-TR8 and TL9-TR5, formed early before the majority of the other tertiary contacts were fully formed (Fig. 4). The reason for the more pronounced involvement of TL2-TR8 and TL9-TR5 in misfolding is that topological entanglement can easily occur when contacts in peripheral regions form first. We surmise that this mechanism is common to the folding of larger ribozymes.

Very few experiments have reported on the details of the misfolded structure due to the difficulty in distinguishing heterogeneous and transient structures. In *Tetrahymena* ribozyme, it was suggested that the misfolded state has a similar topology to the native state, but with less non-native pairing [65]. The misfolded state is mostly stabilized by native-like interactions, but there is “strand-crossing”, by which the RNA conformation is trapped and unable to recover the native structure. Our results show that this is also the case in the smaller and faster-folding *Azoarcus* ribozyme. The metastable states found here consist of correct base pairs and tertiary interactions, which imply that they are native-like topological kinetic traps.

Recently, two groups independently reported cryo-EM structures of the topologically trapped state of the *Tetrahymena* group I intron [44, 45], that shares much in common with the architecture of *Azoarcus*. In both these structures, as predicted here, the J8/7 strand of the central core regions is in an incorrect position relative to the P3 and P7 helices. In Fig. 7, a comparison of these structures with our simulated misfolded structure (J8/7) shows remarkable agreement in the strand arrangement. We surmise that our simulations, conducted without any prior knowledge of the topologically trapped states, predict the major structures of the metastable states that are populated during *Azoarcus* ribozyme folding. Our simulations allow us to trace the process of this misfolding back to the intermediate state (Fig. 7b), thus establishing a kinetic basis for their formation. Compared to the intermediate state in the correct folding pathway (Fig. 7a), we now see how the subtle difference in the folding trajectory of the J8/7 strand ultimately leads to the native and topologically trapped native-like states, both of which are stabilized by the peripheral contacts in a similar way.

### Role of monovalent ions

*Azoarcus* ribozyme folds into a compact structure at high concentrations of monovalent ions even in the absence of Mg^2+^ [66]. However, Mg^2+^ is essential for splicing activity [47]. Nevertheless, both previous experiments [66] and the recent empirical observation [67, 68] that roughly 1 mM Mg^2+^ is equivalent to 80 mM of monovalent cations raise the possibility that high monovalent cations could substitute for Mg^2+^. Therefore, it is interesting to pose the following question. Are the pathways explored by the ribozyme at high monovalent concentrations or when folding is driven by Mg^2+^ equivalent? This question can only be answered using simulations of the kind reported here. However, such simulations are demanding because large system sizes are needed to obtain reliable results. Despite the difficulties, this would be a problem worth investigating in the future.

## Data Availability

The model structures are available in the online supplementary material. The simulation code and parameter files were deposited in Zenodo https://doi.org/10.5281/zenodo.8246270.

## Supplementary Data

Supplementary Data are available at NAR online.

## ACKNOWLEDGMENTS

NH is grateful to Natalia Denesyuk for insightful discussions during the early stage of this study. We appreciate useful discussions with Sarah Woodson and Rick Russell. We thank Anne Bowen in the Texas Advanced Computing Center (TACC) at the University of Texas at Austin for rendering the simulation movies. We acknowledge the TACC for providing computational resources.

