## Supplementary Data for "Watching ion-driven kinetics of ribozyme folding and misfolding caused by energetic and topological frustration one molecule at a time"

(Dated: Aug 21, 2023)

### I. SUPPLEMENTARY METHODS

#### I.1. Coarse-grained RNA model with explicit ions

We used the Three-Interaction-Site (TIS) RNA model [1, 2] in which each nucleotide is modeled using three spherical particles, corresponding to sugar (S), base (B), and phosphate (P) [1] (Figure S1). All the ions, $Mg^{2+}$, $K^{+}$, and $Cl^{-}$, including those in the buffer, are explicitly treated. The solvent (water) is treated implicitly using a temperature-dependent dielectric constant. The TIS force field is written as,

$$U_{\mathrm{TIS}} = U_{\mathrm{bond}} + U_{\mathrm{angle}} + U_{\mathrm{EV}} + U_{\mathrm{HB}} + U_{\mathrm{ST}} + U_{\mathrm{ele}}. \tag{1}$$

The first two terms, $U_{\mathrm{bond}}$ and $U_{\mathrm{angle}}$ ensure the connectivity of the ribose backbone and bases with appropriate rigidity. They are modeled using harmonic potentials,

$$U_{\mathrm{bond}} = \sum_{\mathrm{bonds}} k_b \left(\rho - \rho_0\right)^2 \tag{2}$$

$$U_{\mathrm{angle}} = \sum_{\mathrm{angles}} k_a \left(\theta - \theta_0\right)^2, \tag{3}$$

where the reference values, $\rho_0$ and $\theta_0$, are taken from the ideal A-form RNA helix in a sequence-dependent manner.

*Excluded Volume Interactions:* The term $U_{\mathrm{EV}}$ in equation (1) accounts for volume exclusion that prevents overlap between the CG sites. We used a modified version of the Lennard-Jones type potential,

$$U_{\mathrm{EV}}(r) = \begin{cases} \sum_{(i+1<j)} \varepsilon_{ij} \left[ \left( \frac{a}{r+a-D_{ij}} \right)^{12} - 2 \left( \frac{a}{r+a-D_{ij}} \right)^{6} + 1 \right] & (r \leq D_{ij}) \\ 0 & (r > D_{ij}) \end{cases}. \tag{4}$$

The exclusion distance is $D_{ij} = R_i + R_j$. The energy scale, $\varepsilon_{ij} = \sqrt{\varepsilon_i \varepsilon_j}$, is calculated using the standard combining rule. The values of $R_i$ and $\varepsilon_i$ are tabulated in Table S1. The choice of the distance $a = 0.16$ nm ensures that $U_{\mathrm{EV}}$ becomes a standard Lennard-Jones potential for a pair of interaction sites with the smallest particle type, $Mg^{2+}$ ions.

*

†

*Hydrogen Bond Potential:* We consider hydrogen bonding interactions between bases for all possible canonical base pairs (any G-C, A-U, or G-U base pair can form during the folding process), as well as tertiary hydrogen bonds that are found in the crystal structure (PDB 1U6B [3]). Tertiary hydrogen bonds can occur between any CG sites, whereas canonical base pair is formed only between two bases. Hydrogen bond potential is given by,

$$U_{\text{HB}} = \sum_{\text{H-bonds}} U^0_{\text{HB}} \exp\left(-u_1\right) \tag{5}$$

where

$$u_1 = \begin{pmatrix} k_r(r-r_0)^2 + k_\theta(\theta_1-\theta_{1,0})^2 + k_\theta(\theta_2-\theta_{2,0})^2 \\ +k_\psi(\psi-\psi_0)^2 + k_{\psi1}(\psi_1-\psi_{1,0})^2 + k_{\psi2}(\psi_2-\psi_{2,0})^2 \end{pmatrix}, \tag{6}$$

is a penalty for the deviation from the ideal hydrogen bonding configuration that is represented by the subscript of 0. The ideal configuration is taken from the A-form RNA double helix for the canonical base pairs, and from the crystal structure of the ribozyme (PDB 1U6B [4]) for tertiary hydrogen bonds. The list of tertiary hydrogen bondings was generated by the WHAT-IF server [5].

*Base Stacking Potentials:* Stacking interactions consist of two types, $U_{\text{ST}} = U_{\text{SST}} + U_{\text{TST}}$. The first type, $U_{\text{SST}}$, is the sum of secondary base stacking that are interactions between all consecutive nucleotides, $i$ and $i+1$, regardless of whether they are involved in stacking interactions in the crystal structure. Tertiary stacking interaction, $U_{\text{TST}}$, is the sum of the base-stacking interactions between non-consecutive nucleotides, $i$ and $j$ ($|i-j|>1$), found in the crystal structure. The list of tertiary base-stacking interactions for *Azoarcus* ribozyme is given in Table S1 of Ref. [2]. The total stacking-interaction potential is given by,

$$U_{\text{ST}} = \sum_{\text{secondary}} U^0_{\text{SST}} \left(1+u_2\right)^{-1} + \sum_{\text{tertiary}} U^0_{\text{TST}} \left(1+u_1\right)^{-1}, \tag{7}$$

where $U^0_{SST}$ and $U^0_{TST}$ are stability parameters for secondary- and tertiary-stacking interactions, respectively. For secondary stacking, the penalty term is,

$$u_2 = k_r(r-r_0)^2 + k_\phi(\phi_1-\phi_{1,0})^2 + k_\phi(\phi_2-\phi_{2,0})^2. \tag{8}$$

The value of $u_1$ for tertiary stacking is the same form as in equation (6) for the hydrogen-bonding interactions.

*Electrostatic Potential:* The last term in equation (1) is the Coulomb potential for the electrostatic interactions,

$$U_{\text{ele}} = \frac{e^2}{4\pi\varepsilon_0\varepsilon} \sum_{i<j} \frac{Q_i Q_j}{r_{ij}} \tag{9}$$

where $e$ is the elementary charge, $\varepsilon_0$ is the vacuum permittivity, and $Q$ is the charge of each ion type. We used the experimentally-determined temperature-dependent dielectric constant [6],

$$\varepsilon(T_c) = 87.74 - 0.4008T_c + 9.398\times10^{-4}T_c^2 - 1.41\times10^{-6}T_c^3, \tag{10}$$

where $T_c$ is the simulation temperature in degrees Celsius. The ions interact with each other and with the RNA via the excluded volume and electrostatic interactions ($U_{\text{EV}}$ and $U_{\text{ele}}$), whereas the other four terms account for interactions within the RNA molecule.

In the previous study [2, 7], the parameters in the TIS force field were optimized so that the model reproduced the thermodynamics of dinucleotides, several types of hairpins, and an RNA pseudoknot. The model with the optimized parameters also quantitatively reproduced the measured equilibrium properties of *Azoarcus* ribozyme, such as the equilibrium $R_{\rm g}$ as a function of $Mg^{2+}$ concentrations, and $Mg^{2+}$ binding sites [2, 8]. In addition, we obtained excellent agreements with experiments for heat capacities of pseudoknots and hairpins [2, 7]. These successful applications for calculating the equilibrium properties of RNA molecules set the stage for probing the more difficult problem of calculating the folding kinetics of a large ribozyme.

Note that we used a variant of the TIS model [2], not the one published later [9]. In the 2019 model [9], we developed a theory to treat monovalent ions implicitly whereas the divalent ions were treated explicitly. Although the latter model is advantageous in terms of computational cost due to the absence of explicit monovalent ions, it was essential for us to use the explicit representations of both divalent and monovalent ions in the current study. This allowed us to directly quantify the condensation and binding of both monovalent and divalent ions (Figure 6a), and to vividly capture the replacement of $K^+$ ions by $Mg^{2+}$. This would not be possible using the implicit monovalent ion model.

#### I.2. Simulation protocols

In order to generate the folding trajectories, we performed Brownian dynamics simulations. The equation of motion is [10],

$$\dot{\boldsymbol{x}} = -\frac{1}{\gamma}\frac{\partial U_{\rm TIS}}{\partial \boldsymbol{x}} + \boldsymbol{\Gamma}. \tag{11}$$

where $\boldsymbol{x}$ is a coordinate, and $\boldsymbol{\Gamma}$ is a Gaussian random force that satisfies the fluctuation-dissipation relation given by $\langle \boldsymbol{\Gamma}_i(t)\boldsymbol{\Gamma}_j(t')\rangle = 6\gamma k_{\rm B}T\delta(t-t')\delta_{ij}$. The friction coefficient follows the Stokes-Einstein relation, $\gamma = 6\pi\eta R$, where $\eta = 8.9\times10^{-4}$ Pa·s is the viscosity of water, and $R$ is appropriate radius for each coarse-grained site. We set values of $R$ for the RNA sites and ions according to previous studies [11, 12].

The simulation time was mapped to the real time by relating the microscopic time scale $\tau_L = \sqrt{ma^2/\epsilon} \sim 2$ ps and the characteristic time of the Brownian dynamics, $\tau_H = \frac{6\pi\eta a^3}{k_B T} \approx 300$ ps, where the typical energy scale $\epsilon \sim 1$ kcal/mol and the length scale $a \sim 0.4$ nm [13, 14]. We previously demonstrated that this way of predicting kinetics using the TIS model provides *quantitative* agreement for the time scale of RNA folding for hairpin and pseudoknot RNA molecules [15–18].

For each initial conformation, taken from the unfolded ensemble generated as described above, $Mg^{2+}$ and $Cl^-$ ions corresponding to 5 mM $MgCl_2$ were added before initiating the folding simulations. The additional ions were randomly distributed in the cubic box, keeping them at least 0.5 nm from the preexisting ions and the ribozyme. The folding simulations were continued until the RNA reached the native conformation, or the time steps reached $10^{10}$ corresponding to 30 ms, which is the maximum simulation time in all our trajectories. If *Azoarcus* ribozyme did not fold within the maximum simulation time, then it is considered to be kinetically trapped, a phenomenon that frequently occurs in *in vitro* experiments.

The coordinates of the RNA and ions were recorded every 50,000 time steps. We assessed if RNA is folded within the time scale of simulations using the following criterion: Root mean square deviation (RMSD) from the PDB structure was computed every frame. The RMSD is then averaged over a time window of 1,000 frames. If the window average of RMSD is smaller than 0.6 nm, the trajectory is deemed to reach the folded state. In a few trajectories, we found unphysical chain crossing when the length of the pseudo-covalent bond between sugar and phosphate becomes comparable to the excluded distance between them. Although this rarely occurred, it should not be dismissed because such crossing could affect the folding kinetics. Thus, when a chain crossing occurred in a trajectory, we restarted the simulation with a different random number seed to ensure the backbone of RNA does not cross each other.

#### I.3. Analyses

**Finding major misfolded conformations:** The clustering analysis was performed as described in Models and Methods in the main text. The result of the clustering analysis is shown in Figure S7 as a dendrogram along with structures of cluster centers. The compact-structural ensemble is grouped into five parent clusters that are designated as cluster identification (ID) 1 through 5 in the dendrogram. From a visual inspection of the structures, we identified the five clusters as follows: Cluster 1 is Misfolded (P2) conformation (see the main text), Cluster 2 belongs to the native-like ensemble, Clusters 3 and 5 are both Misfolded (J8/7) conformation, and Cluster 4 is another minor misfolded state that appeared in only one trajectory. Although structures in Clusters 3 and 5 are classified into two distinct clusters, they have the same chain topology and tertiary contacts, except that TL9-TR5 contact is formed in Cluster 3, whereas it is disrupted in Cluster 5. For sub-clusters 1a and 1b, 2a and 2b, and so on, we confirmed that they represent minor variations of the root clusters. This method provides an unbiased way to classify the misfolded structures, which are, except in rare cases (Cluster 1), difficult to observe in experiments.

**Calculation of $\mathbf{Mg^{2+}}$ binding rate:** The mean first passage time (MFPT) of $Mg^{2+}$ to bind to each phosphate site was calculated as follows. The phosphate associated with nucleotide site $i$ in a trajectory $j$ is considered to be bound by $Mg^{2+}$ if the distance to the closest $Mg^{2+}$ ion is shorter than the cutoff distance in 10 consecutive time frames (1.5 $\mu s$). The time at which the binding occurs for the first time, $t_{i,j}$, was recorded as the first passage time of the trajectory. The MFPT for the $i^{\text{th}}$ nucleotide was calculated as,

$$\tau_i = \frac{1}{N}\sum_{j}^{N} t_{i,j}, \tag{12}$$

where $N$ is the number of trajectories. The $Mg^{2+}$ binding rate of the $i^{\text{th}}$ nucleotide is,

$$k_b(i) = \frac{1}{\tau_i}. \tag{13}$$

Note that, for some nucleotide sites in certain trajectories, $Mg^{2+}$ ions do not bind tightly within the simulation time $T_s$. In such cases, we assign $t_{i,j} = T_s$ in equation (12). In that sense, $k_b$ is an estimate of the upper bound of the binding rate from the simulations because the true first passage time ($t_{i,j}$) would be larger in such trajectories.

**Definition of the folded helix, tertiary interactions, and non-native helix:** The fraction of helix formation (Figure 5e) was calculated as the fraction of base pairs formed in the helix at a specific time. A base pair was assumed to be formed if the magnitude of the hydrogen-bonding energy (equation 5) was more favorable than the thermal energy, $k_{\mathrm{B}}T$. Similarly, the formation of key tertiary interactions (Figure 1 and 5f) was assessed by comparing the tertiary hydrogen-bonding or stacking energy to the thermal energy.

In order to find the major mispaired helices, *i.e.* double-stranded helices consisting of non-native base pairs shown in Figure S8, we first made a list of non-native base pairs, that contained all possible canonical base pairs (A-U, G-C, and G-U) that are not present in the crystal structure. Then we counted the frequencies of each non-native base pair over all the trajectories. Lastly, we sorted the list by the frequencies of their formation. The major mispaired helices were identified from combinations of base pairs that formed most frequently, which appeared at the top of the sorted list. The fractions of mispaired helices (Figure S21) were calculated by checking whether the mispaired helices are formed or not at each time step in each trajectory. If two or more non-native base pairs formed, the helix was deemed to be formed.

### II. SUPPLEMENTARY DISCUSSION

**Folding kinetics of helices:** To investigate the details of the folding process, we focussed on the formation of individual secondary structural elements. We calculated the time-dependent folded fraction of helices using the trajectories which reached the native structure (Figure 5e). There are eight helices, P2 through P9 in the *Azoarcus* ribozyme (Figure 1a). At $t = 0$ and in the absence of $Mg^{2+}$, helices P2, P4, P5, and P8 are stably folded, which are also found in equilibrium simulations (Figure 5e). The presence of stable helix regions, including the partial formation of P6 and P9 (see the following), is consistent with experimental results [19]. Consequently, the fractions of these four helices start from high ($> 80\%$) at $t = 0$, and gradually increase throughout the folding process. A part of the P9 helix can be sectionalized as P9.0, which is not stable in the absence of $Mg^{2+}$. Due to the unstable part, the fraction of P9 is about ∼0.7 in the early stage of folding (purple line in Figure 5). Nevertheless, the entire P9, including P9.0, formed rapidly in $t < 0.1\,\text{ms}$ upon titrating with 5 mM $Mg^{2+}$. Helix P6 also has a sub-region, P6a, which is more stable than the rest. The six base-pairs at the tip of P6 are stable even in the absence of $Mg^{2+}$, contributing to about 40% formation of the helix in the initial stage (light green in Figure 5). The other base pairs in P6 fold on a longer time scale. Formation of the three base pairs involved in the Tertiary-Helix (TH, see 1b) is particularly slow; it folds in the final stages, $t > 10\,\text{ms}$.

Even though some parts of P6 and P9 fold more slowly than others, helices P2, P4, P5, P6, P8, and P9 are essentially simple stem-loop structures, with the two strands being located close to each other along the sequence. Therefore, we surmise that there is a negligible entropic barrier for their formation so that they fold rapidly (or even form, to a great extent, in the absence of $Mg^{2+}$). Helices P3 and P7 exhibit particularly slow folding kinetics (light and dark blues in Figure 5). The two strands of P3 and P7 helices are far along the sequence, and thus it takes more time to search for their counterparts.

**A minor topological trap by incorrect positioning of the P2 helix:** In addition to the major topologically trapped state (Misfold J8/7), described in the main text, we found another minor topological misfolding due to incorrect positioning of the P2 helix domain. In this Misfold P2, one of the P3 strands leading to the P2 domain is positioned in the opposite direction relative to its native position when the TL9-TR5 tertiary interaction forms, thus fixing the P2 domain on the wrong side (Figure S9). The space between the two pairs of coaxial helix domains P4-P5-P6 and P7-P8-P9 is too narrow for the P2 helix to move to its normal position on the opposite side unless TL9-TR5 dissociates.

By searching through all the folding trajectories using the representative structure from the clustering analysis, we found that the misfolded state occurred in only three out of 95 trajectories. In two of the three trajectories, the TL9-TR5 interaction persisted after the formation of the P2 misfolded state, and therefore the ribozyme remained trapped throughout the simulation (Figure S10). In the other trajectory, the TL9-TR5 interaction was eventually disrupted by thermal fluctuations and the P2 misfolded state was resolved (Figure S11). However, the ribozyme did not fold to the native structure within the simulation time. Although it is difficult to deduce the exact mechanism by which this misfolded state occurs from only three trajectories, what is common in these trajectories is the partial unfolding of the P6 helix and the relatively early formation of the G site (Figs. S10 and S11).

**TH formation and $Mg^{2+}$ binding to U51 and U126:** Triple Helix (TH) is a structural element that folds in the early stage following Stack Exchange (Figure 5f). As shown in Figure S18, nucleotides U51 and U126 bind $Mg^{2+}$ ions upon folding of the TH, which is consistent with the observation that they are coordinated by two $Mg^{2+}$ ions in the crystal structure. The correlation coefficients of the first passage times are relatively low, $\rho = 0.58$ and 0.73 for U51 and U126, respectively. The scatter plots in Figure S18(c, d) show that, in some trajectories, TH forms earlier than $Mg^{2+}$ binding to U51 and U126, indicating that $Mg^{2+}$ binding to these nucleotides are not a strict requirement for TH to form. This is consistent with the finding that TH forms at high probability (≈0.6) even at submillimolar $Mg^{2+}$ concentration at equilibrium [2].

**Correlation between SE formation and $Mg^{2+}$ binding to G41, G138 and A167-A168:** The Stack Exchange junction (SE) is the coaxial stacking between P3 and P8 helices, whose formation facilitates the folding of the P3 pseudoknot. Figure S17 shows that the formation of SE temporally correlates with the $Mg^{2+}$ binding at a cavity formed by G41, G138, and A167-A168. The correlation coefficients are all above 0.9. Based on the result of equilibrium simulations [2], it was conjectured that $Mg^{2+}$ ions bind this region in a diffusive manner, which makes it difficult to resolve $Mg^{2+}$ densities in the crystal structures. The results of our kinetics simulations also support the finding that $Mg^{2+}$ binding events to these nucleotides are important for the formation of the SE motif.

**Correlation between $Mg^{2+}$ binding and formations of peripheral interactions:** Two peripheral tertiary motifs, TL2-TR8 and TL9-TR5, are also stabilized by $Mg^{2+}$ ions. As shown in Figure S19 and S20, both peripheral motifs have two or three nucleotides having high temporal correlations between $Mg^{2+}$ binding and formations of the motif. There is, however, a qualitative difference between the two cases. In the TL2-TR8 case, the tertiary interaction may form earlier than the time for specific $Mg^{2+}$ binding events, although they are at the same time in the majority of the trajectories (Figure S19 a-b). We did not observe any trajectory in which $Mg^{2+}$ bound before the motif was formed. In contrast, $Mg^{2+}$ binding to TL9-TR5 motif can be earlier or later than the formation of the interaction (Figure S20 a-c). These events are temporally correlated. The Pearson correlation coefficients of the first passage times are higher for TL9-TR5 ($\rho = 0.79$, 0.82, 0.97) than for TL2-TR8 ($\rho = 0.58$ and 0.73). This may be related to the subtle difference in their stability observed in equilibrium simulations, that is, TL9-TR5 unfolded rapidly at low $Mg^{2+}$ concentrations (1 mM) whereas TL2-TR8 was still partially folded ($\sim 30\%$ population) [2].

TABLE S1. **Parameters for excluded volume and electrostatic interactions.** In the first column, P is phosphate, S is sugar, and A, G, C, and U are the four bases. The charge, $Q_i$, is defined in units of the proton charge. If both interacting sites are RNA sites, we take $R_i + R_j = 0.32$ nm.

| | $R_i$, nm | $\varepsilon$, kcal/mol | $Q_i$ |
|---|---|---|---|
| P | 0.21 | 0.2 | -1 |
| S | 0.29 | 0.2 | 0 |
| A | 0.28 | 0.2 | 0 |
| G | 0.30 | 0.2 | 0 |
| C | 0.27 | 0.2 | 0 |
| U | 0.27 | 0.2 | 0 |
| $Mg^{2+}$ | 0.08 | 0.9 | 2 |
| $Cl^{-}$ | 0.19 | 0.3 | -1 |
| $K^{+}$ | 0.27 | 0.0003 | 1 |

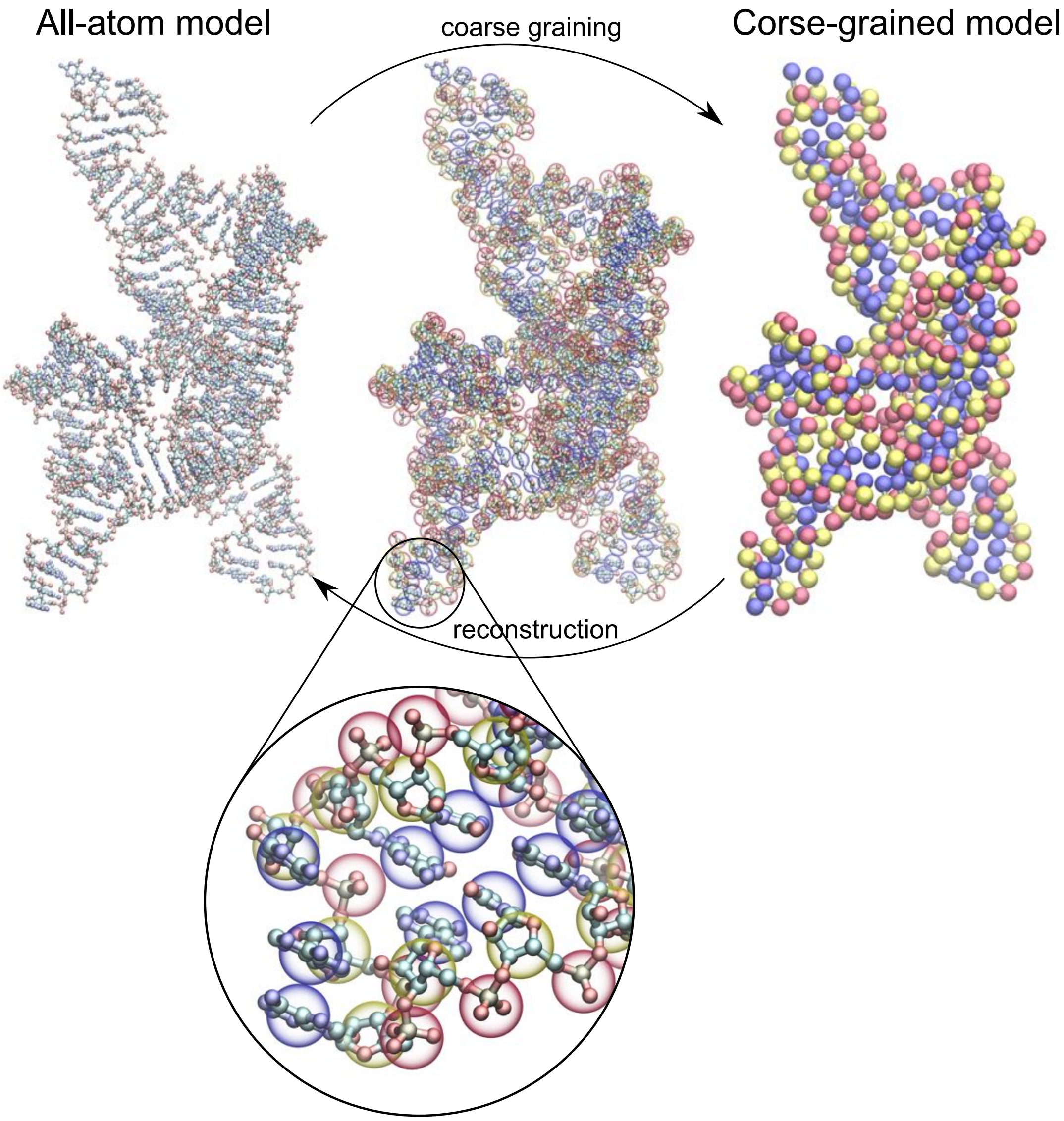


FIG. S1. **Three-Interaction-Site model.** All-atom representation of *Azoarcus* group I intron RNA (left) is compared to the TIS coarse-grained representation (right). In the middle, the two representations are overlaid, and a part of the RNA is magnified at the bottom. The coarse-grained sites are colored in red (phosphate), yellow (sugar), and blue (base).

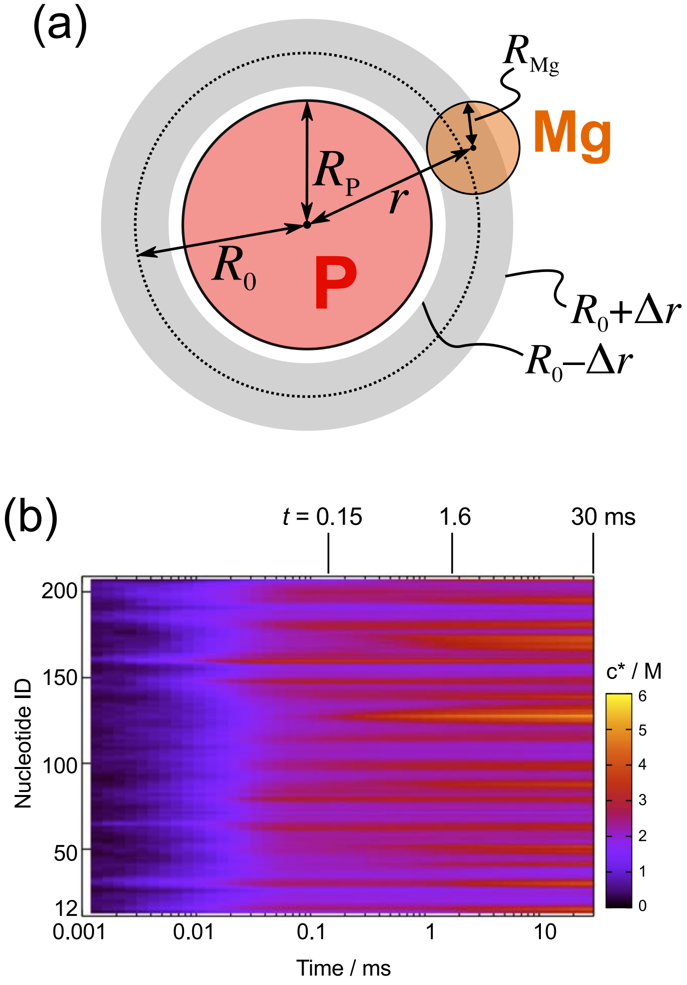


FIG. S2. **Time dependent binding of $Mg^{2+}$.** **(a)** A schematic cross-section of a $Mg^{2+}$ ion bound to a phosphate group (P), used to define the contact $Mg^{2+}$ concentration, $c^*$. We counted the number of $Mg^{2+}$ ions located in the range, $r_0 - \Delta r < r < r_0 + \Delta r$, where $r$ is the distance from the phosphate, $r_0 = R_\mathrm{P} + R_\mathrm{Mg}$ is the sum of the excluded-volume radii, and $\Delta r = 0.15$ nm is a margin. The contact $Mg^{2+}$ concentration is then calculated by dividing the number by the spherical shell volume (shaded in grey). **(b)** Average contact $Mg^{2+}$ concentration at each nucleotide as a function of time. In the main text, $Mg^{2+}$ fingerprints are shown as the profile of $c^*$ at $t = 0.15$, 1.6, 30 ms (Figure 6). The heterogeneity of association is considerable.

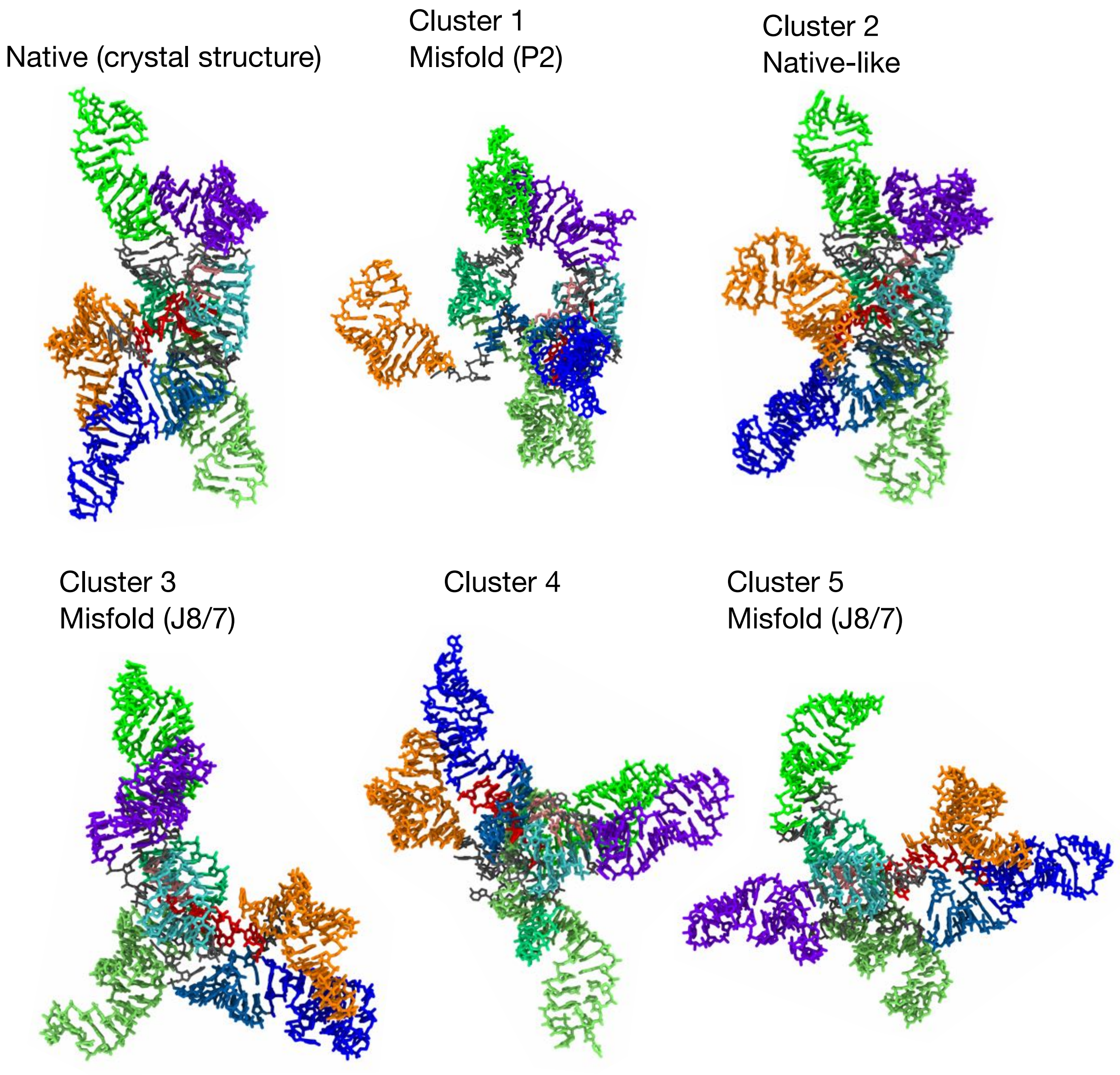


FIG. S3. **Atomistic representations of cluster-center structures.** The same representative misfolded structures as shown in Figure S7 with the native structure (top left). The structures are reconstructed from coarse-grained coordinates using the fragment-assembly approach followed by energy minimization, as described in Methods.

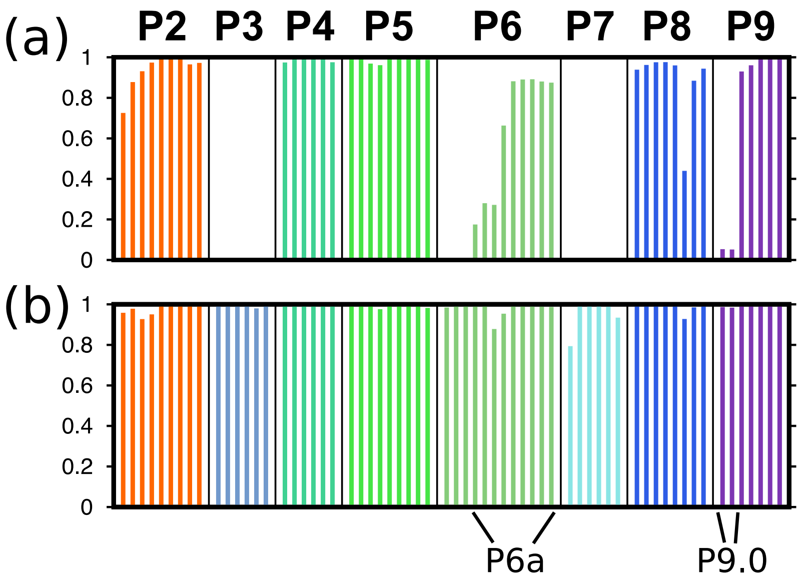


FIG. S4. Probabilities of helix formation for helices P2 to P9 under equilibrium conditions in the (a) absence and (b) presence (5 mM) of $Mg^{2+}$. The labels for helices are shown on top of each block. Each bar represents a base pair consisting of each helix. P6 and P9 are sectioned into parts, P6a and P9.0, which are indicated in the bottom panel (see also the secondary structure in Figure 1a in the main text).

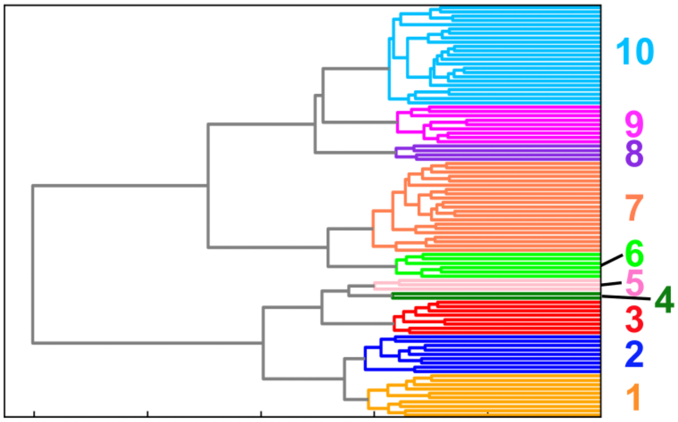


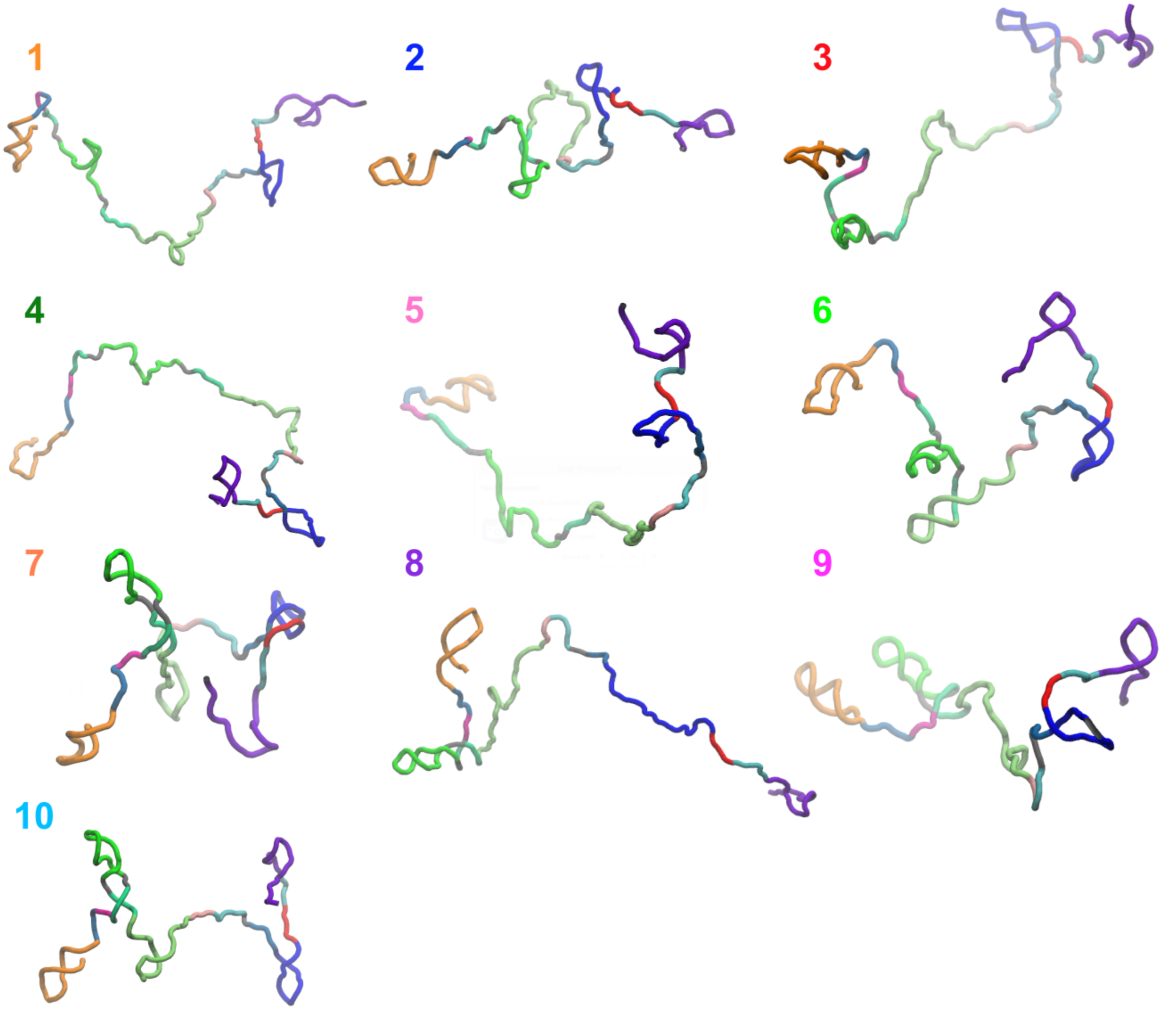


FIG. S5. **Hierarchical clustering of initial structures.** The dendrogram (top) shows the result of clustering analysis for the 95 initial structures with DRID as the distance metric, in the same manner as the clustering of compact structures. Representative structures of the top 10 clusters are shown. The unfolded conformations are structurally diverse.

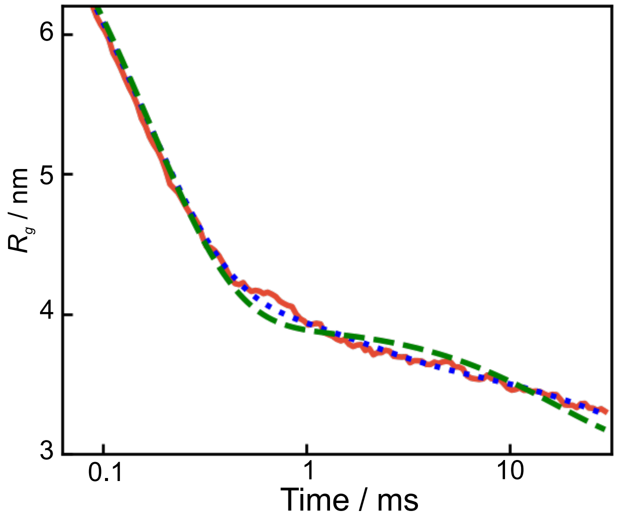


FIG. S6. **Fits of ensemble-averaged radius of gyration ($R_\mathbf{g}$) as a function of time.** The solid red line is $\langle R_\mathrm{g} \rangle$, calculated from all the simulation trajectories. The green dashed line is the best fit using a sum of two exponential functions ($\Phi_1 = 0.81$, $\Phi_2 = 0.19$, with the corresponding time constants, $\tau_1 = 0.17$, $\tau_2 = 17$ ms). The blue dotted line is the fit using three exponential functions discussed in the main text. The sum of two exponential functions does not quantitatively capture the time dependence beyond $\sim$1 ms. There is a noticeable deviation between the red and green curves.

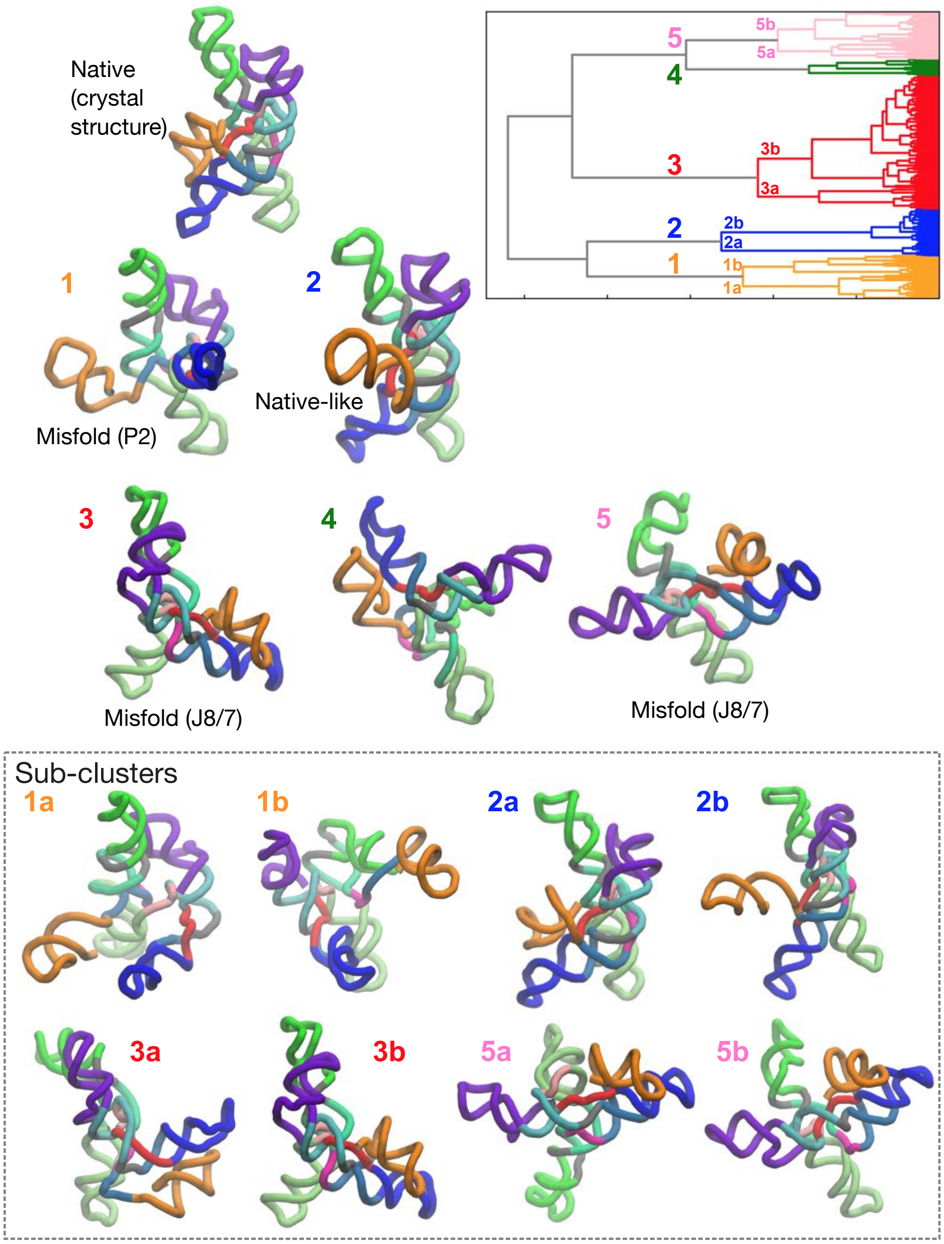


FIG. S7. **Clustering of compact structures from the simulation trajectories.** (Top right) The dendrogram from the hierarchical clustering based on pairwise DRID. Cluster IDs are assigned for several representative clusters. (Top left) The crystal structure of the ribozyme is shown for comparison. (Middle) Structures from the five major cluster centers. Cluster 2 contains the native topology conformations. The other four clusters are misfolded conformations. (Bottom) Structures of sub-clusters of Cluster 1, 2, 3, and 5 are also shown for comparison.

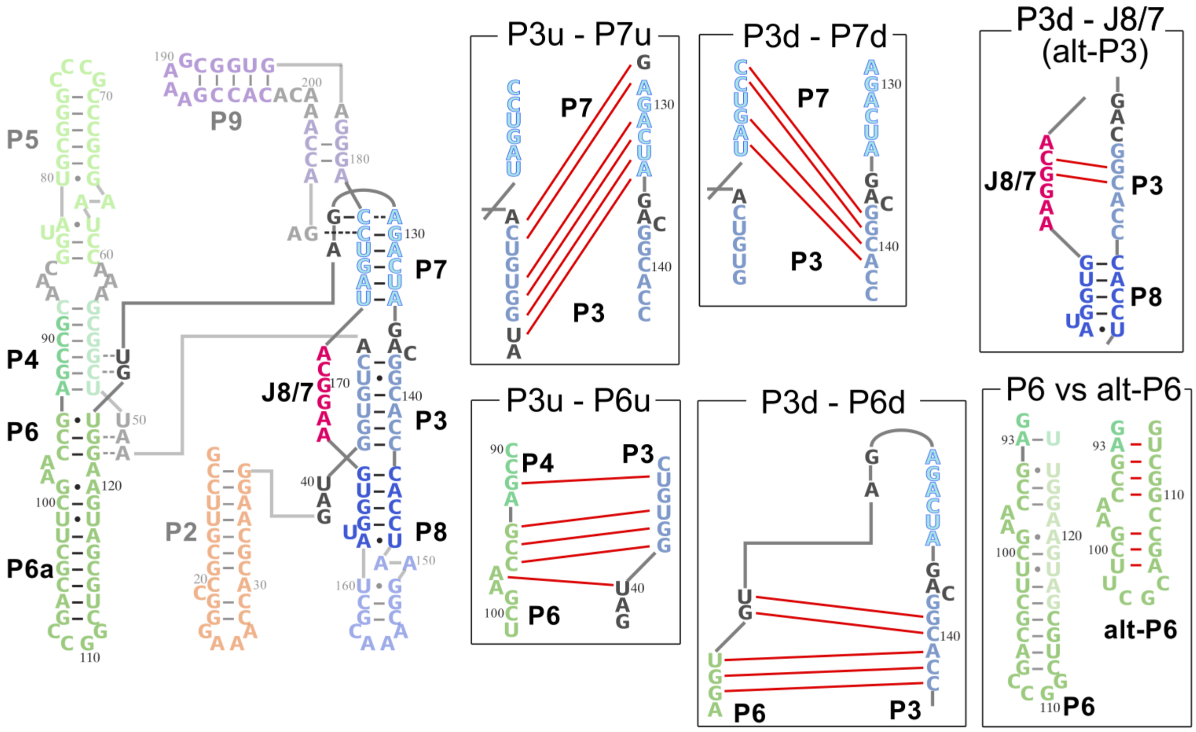


FIG. S8. **Representative mispairing found in the simulations.** The native secondary structure is shown on the left. Six frequently-observed mispairing patterns are shown in rectangles on the right. Each mispairing pattern is named with constituent strands, where each strand is distinguished as either up-stream (u) or down-stream (d) in helices.

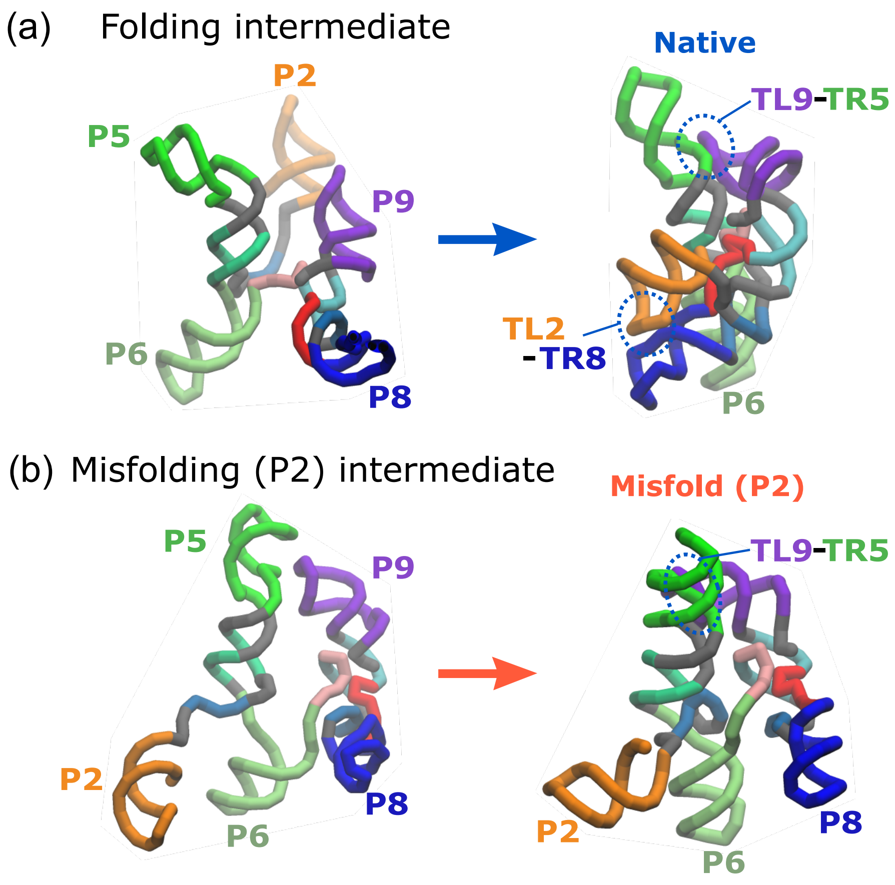


FIG. S9. **Comparison between intermediate states leading to the P2 trapped state and correct folding.** Two intermediate structures are extracted from trajectories (a) folding to the native structure and (b) misfolded (P2) state. The two intermediate states were visualized so that the relative positions of P5, P9 and P6 were approximately the same. In the correct folding (a, left), the P2 domain is positioned behind P5 and P9 helices, whereas in the misfolding (b, left) it comes to the front. Once the TL9-TR5 contact is formed in this situation, the P2 domain is unable to move backwards (b, right).

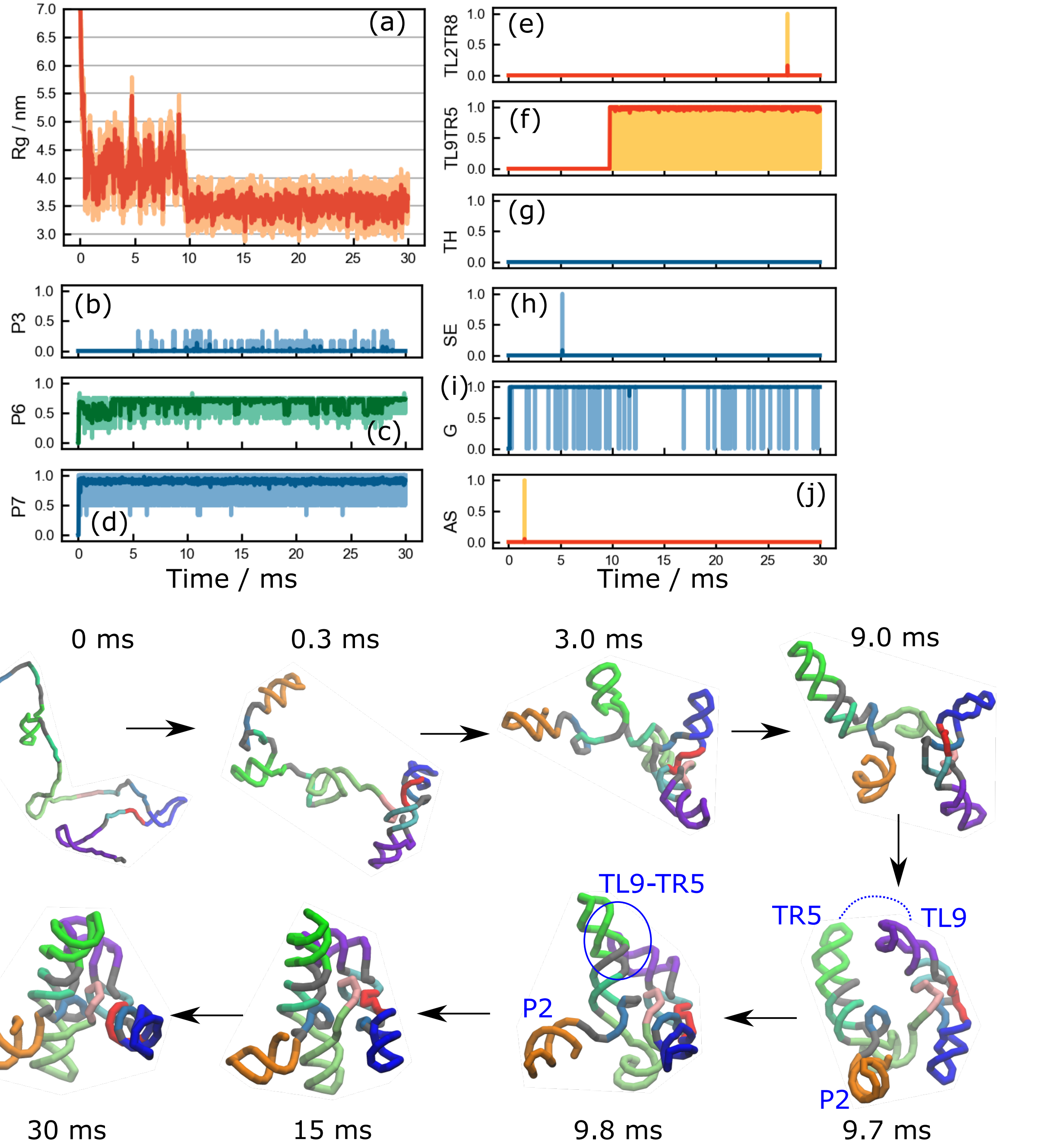


FIG. S10. **Time course of a trajectory kinetically trapped to Misfold (P2) state.** One of the three trajectories in which the misfolded (P2) state was found. In this trajectory, at $t$ = 9.8 ms, the TL9-TR5 tertiary interaction formed when the P2 helix domain was mispositioned. The space between helices P3-P4-P5 (green) and P7-P8-P9 (blue, cyan, purple) is too narrow for the P2 helix to move back to the other side. Therefore the misfolded state remains until the end of the simulation.

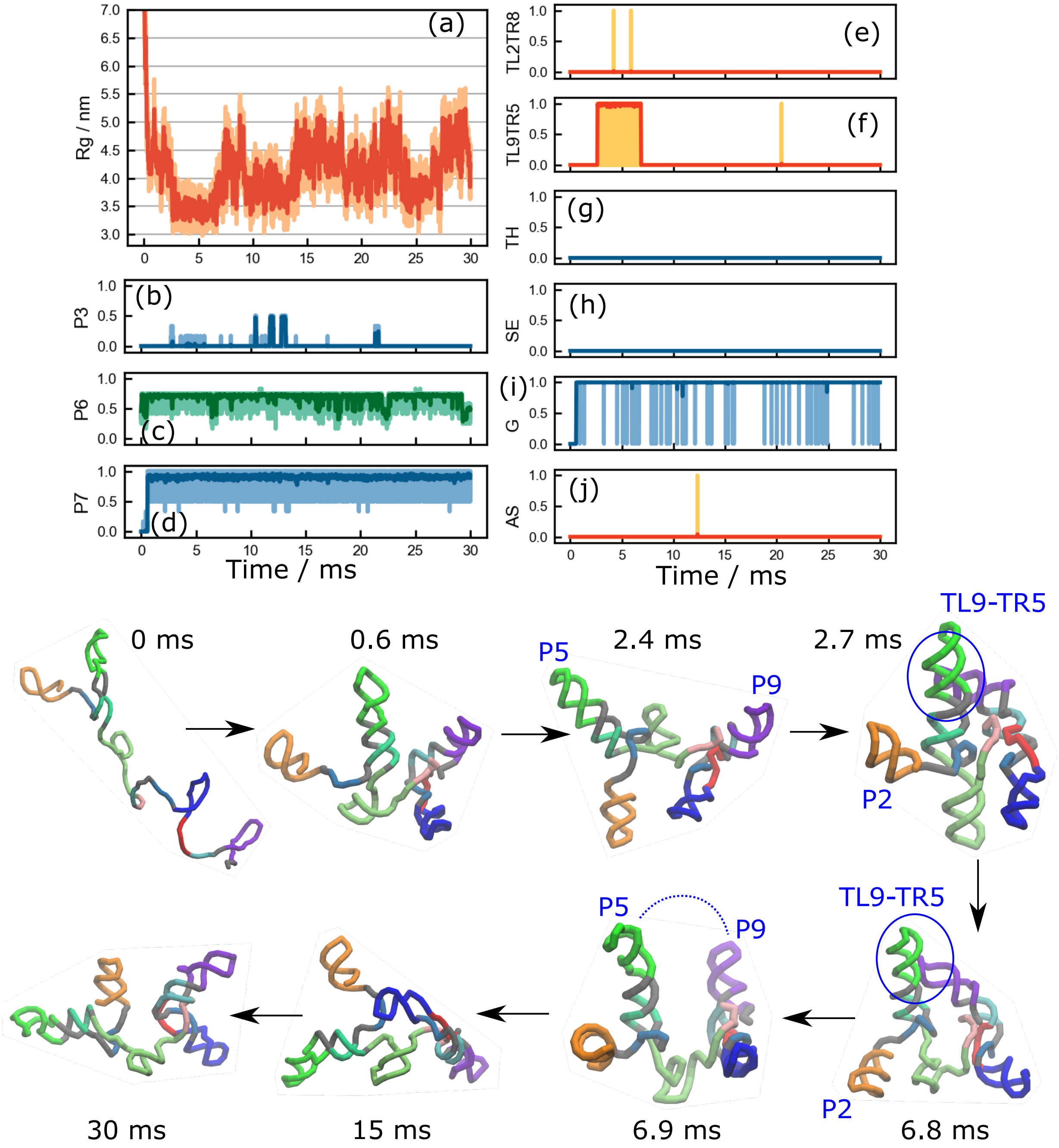


FIG. S11. **Escape from the Misfold (P2) state.** This is the only trajectory where we observed escape from a topological trap. Early in the trajectory ($t$ = 2.7 ms), the TL9-TR5 tertiary contact is formed while the P2 helix domain was in the misfolded position, in the same manner as in Figure S10. The ribozyme remained in the misfolded state until $t \sim 6.8$ ms. Eventually, the TL9-TR5 interaction disengaged by the thermal fluctuation and the misfolded state was resolved. However, folding to the native structure did not occur within the simulation time.

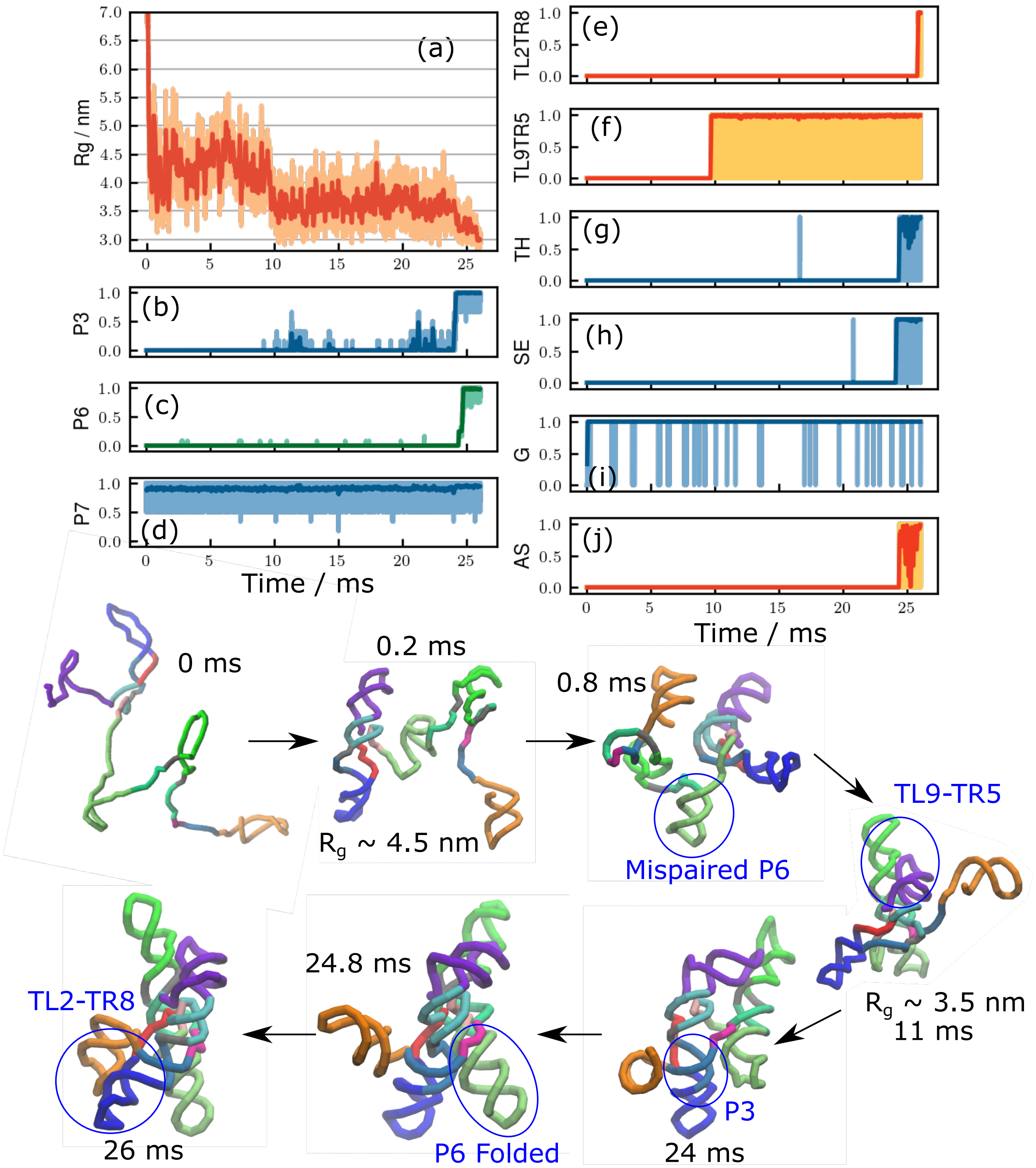


FIG. S12. **Folding trajectory.** A trajectory that reached the folded state in a time scale larger than the one shown in the main text. A mispaired helix in P6 formed early ($t \sim 0.8$ ms), which prevents it from folding to the native state rapidly. Eventually ($t \sim 24$ ms), the misfolded helix was resolved, leading to the correct native fold. It should be pointed out that commitment to rapid specific collapse, defects in collapse but resolvable or trapping for arbitrarily long times in compact misfolded states occur roughly on the collapse time scale.

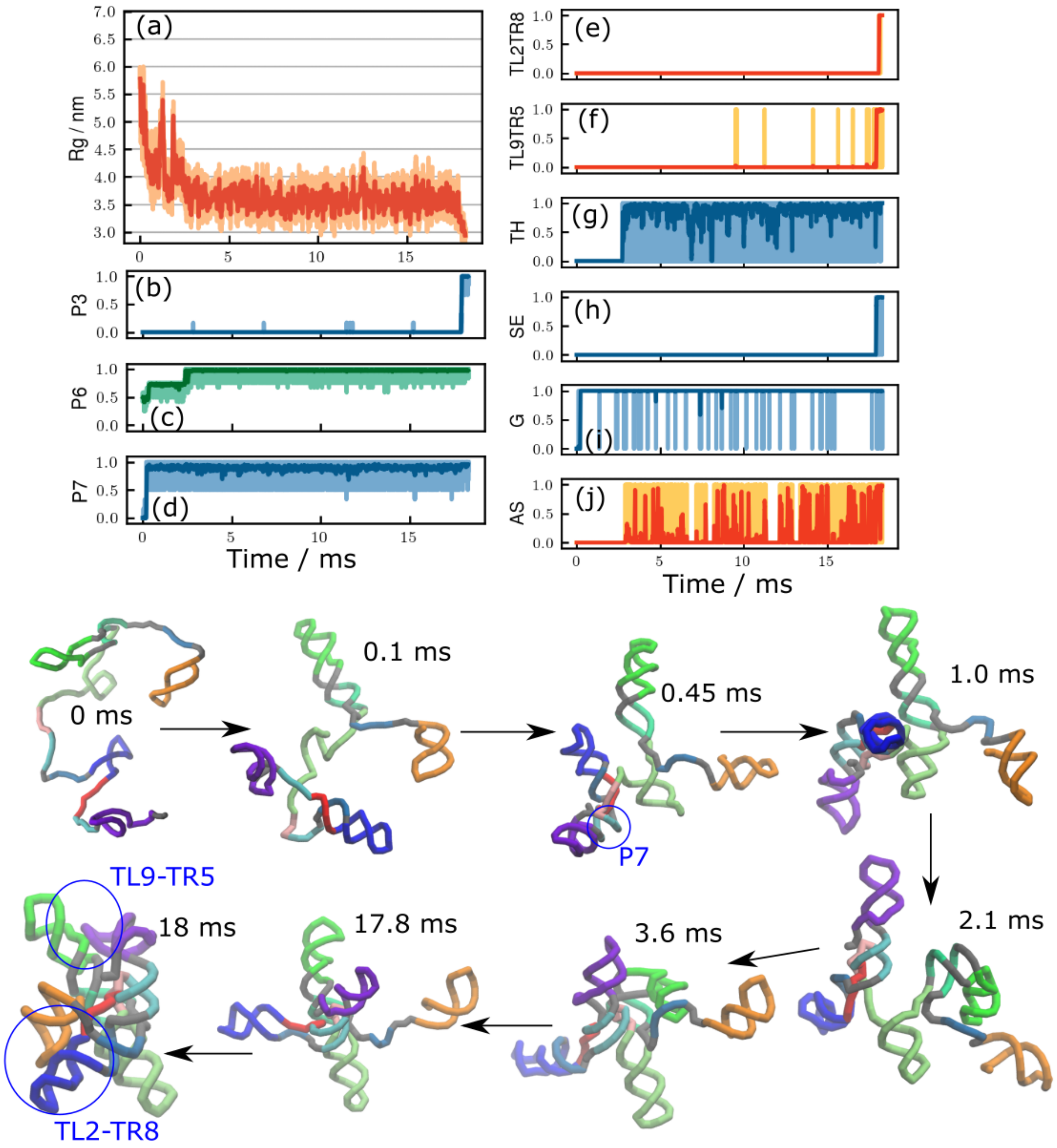


FIG. S13. **Different routes to the folded state.** As with Figure S12, this is another trajectory that reached the folded state in a time scale larger than the one shown in the main text. In this trajectory, helices P6 and P7 folded earlier ($t \sim 2$ ms), followed by the formation of the Triple Helix (TH). Two peripheral interactions, TL2-TR8 and TL9-TR5, needed longer times to search the counterparts. The folding was completed at $t \sim 18$ ms. There is no typical folding trajectory.

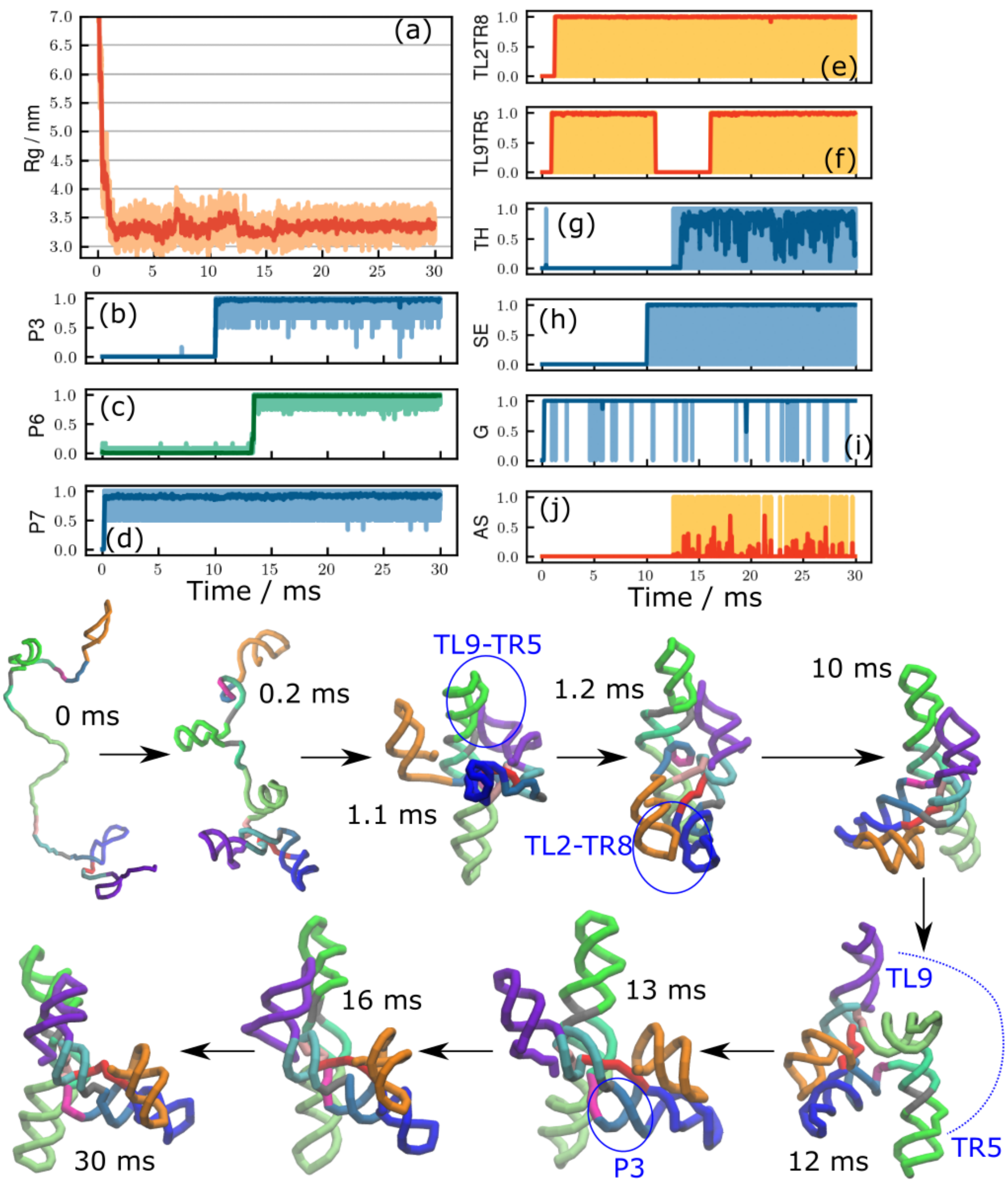


FIG. S14. **Time course of a kinetically trapped trajectory.** In this trajectory (also shown in Figure 4), the mechanism of formation of a persistent misfolded state is revealed. Two key interactions in the peripheral regions, TL2-TR8 and TL9-TR5, formed at times $t < 1$ ms, result in the junction J8/7 with an incorrect topology (See Figure 7b). Because the incorrect chain topology cannot be resolved unless both peripheral interactions unfold, the RNA stays in this misfolded state for the rest of the simulation time. The trapped structure is an example of topological frustration.

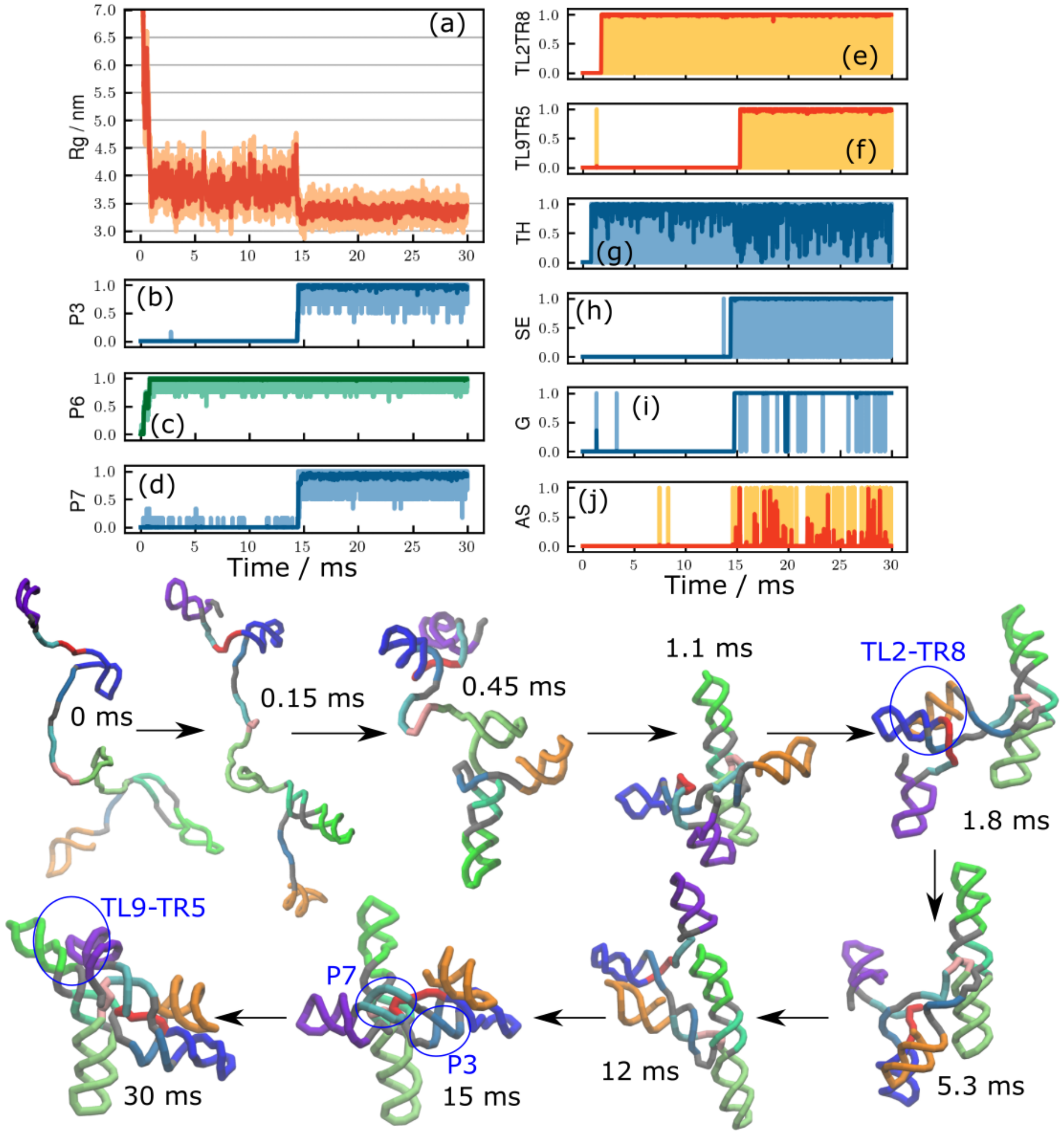


FIG. S15. **Illustrating heterogeneity in the kinetically trapped trajectories.** As with Figure S14, this is another example of a kinetically trapped trajectory, that reached the same misfolded (J8/7) state. In this trajectory, it took a long time ($t \sim 15$ ms) to form a compact intermediate state ($R_{\rm g} \sim 3.5$ nm) mainly because P3 and P7 helices did not form earlier. Also, unlike the previous example (Figure S14), one of the peripheral interactions, TL9-TR5, formed in the later stage.

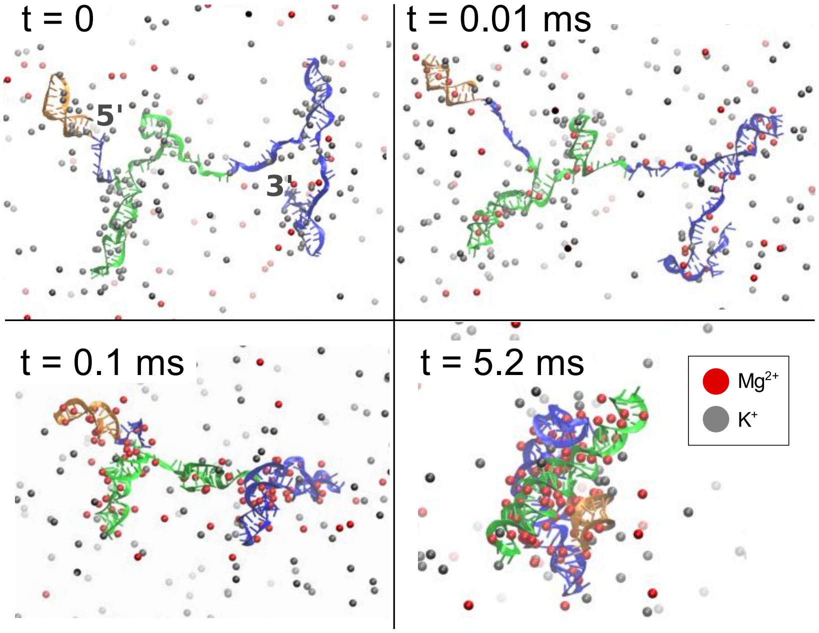


FIG. S16. **Snapshots of ion condensations chosen from a folding trajectory.** The color scheme for ions is shown in the figure. For visual clarity, only cations in the vicinity of the RNA are shown. The size of spheres does not reflect actual volumes in the simulations. Although the majority of the ions near the folded RNA (last panel) are $Mg^{2+}$, there are detectable $K^{+}$ ions that also interact with the ribozyme in a site-specific manner. This implies that monovalent ions, even at low concentrations, could be present in the vicinity of the folded ribozyme.

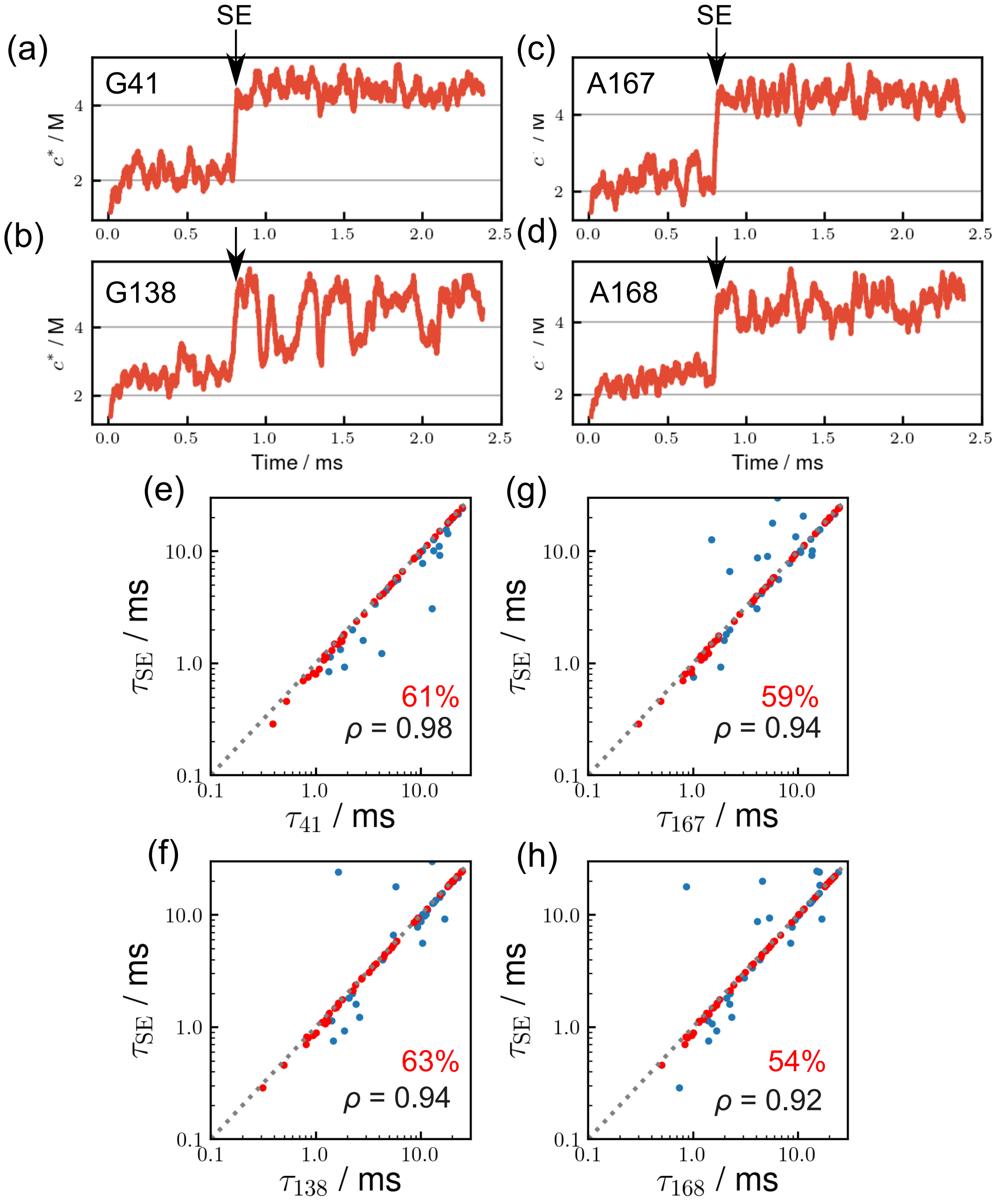


FIG. S17. **Time correlation between $Mg^{2+}$ binding and the formation of Stack Exchange (SE).** (a-d) Trajectories of $Mg^{2+}$ binding, measured in terms of contact concentration, at (a) G41, (b) G138, (c) A167, and (d) A168 taken from the same simulation trajectory as Figure 3. The folding time of SE is indicated by black arrows. (e-h) Scatter plots of the first passage times of $Mg^{2+}$ binding to the same four nucleotides versus the first passage time of SE formation. Cooperative binding of $Mg^{2+}$ to these sites drives SE formation.

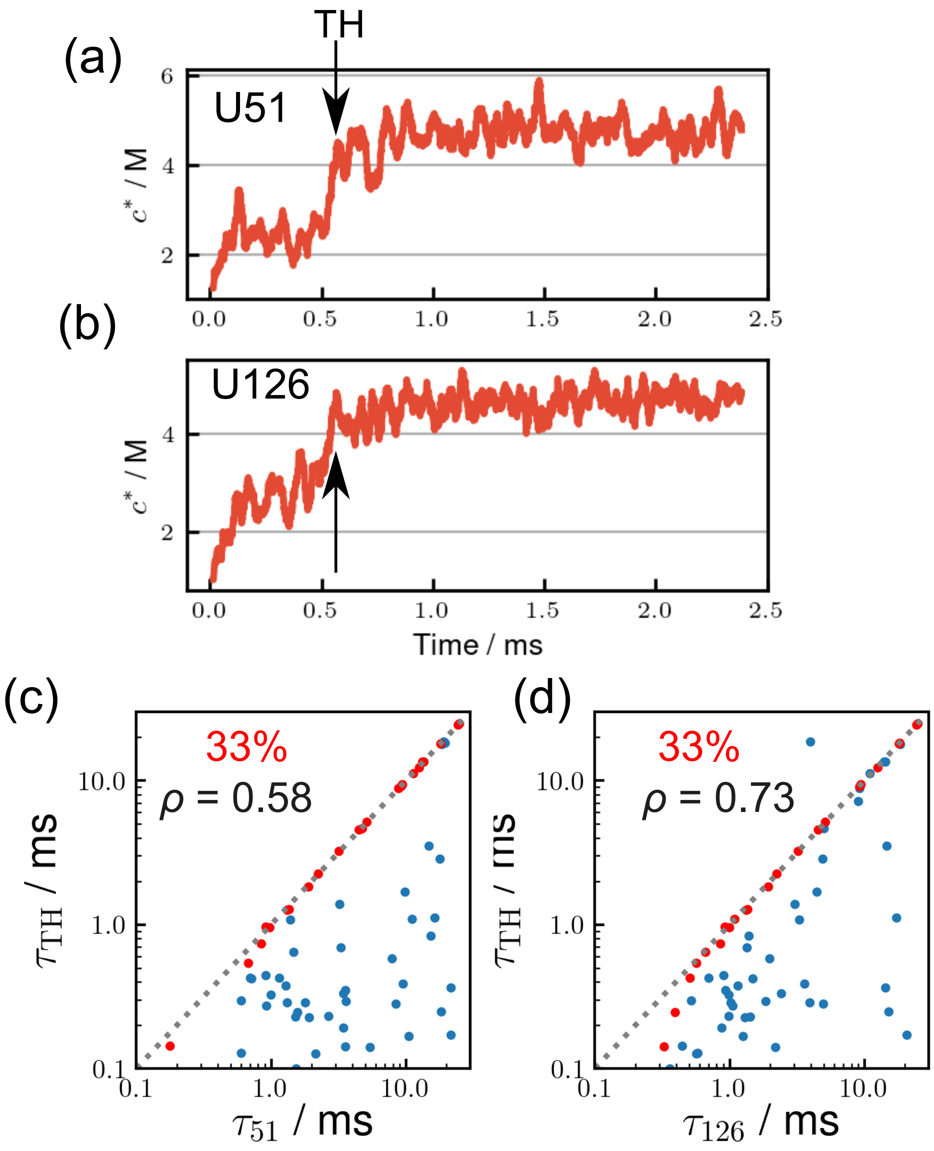


FIG. S18. **$Mg^{2+}$ binding times to U51 and U126 and the formation of Triple Helix (TH). (a, b)** Trajectories of $Mg^{2+}$ binding to (a) U51 and (b) U126 taken from the same simulation trajectory as Figure 3. The folding time of TH is indicated by black arrows. **(c, d)** Scatter plots of the first passage times of $Mg^{2+}$ binding to (c) U51 and (d) U126 versus the first passage time of the TH formation. Unlike the results in Figure S17, there are two modes of $Mg^{2+}$ associations. In one class, TH formation and $Mg^{2+}$ binding are concomitant. There are several trajectories in which $Mg^{2+}$ association occurs after the TH formation (red dots). In this class, there is a broad distribution of $Mg^{2+}$ binding times.

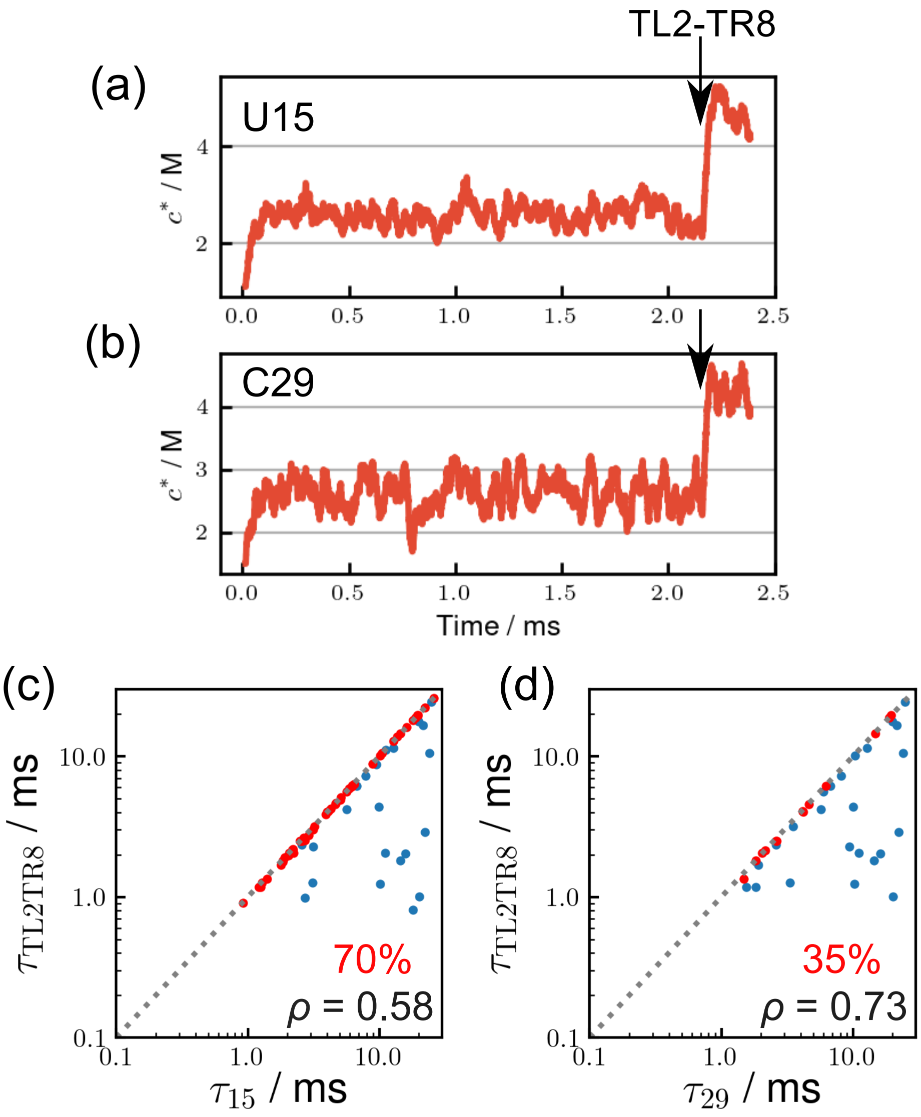


FIG. S19. **Same as Figure S18 except association of $Mg^{2+}$ to U15 and C29 and the formation of TL2-TR8 is shown. (a, b)** Trajectories of $Mg^{2+}$ binding at (a) U15 and (b) C29 taken from the same simulation trajectory as Figure 3. The folding time of TL2-TR8 is indicated by black arrows. **(c, d)** Scatter plots of the first passage times of $Mg^{2+}$ binding to (c) U15 and (d) C29 versus the first passage time of the TL2-TR8 formation. Just as in Figure S17, we find correlated $Mg^{2+}$ binding and TL2-TR8 formation, as well as structure formation followed by $Mg^{2+}$ binding.

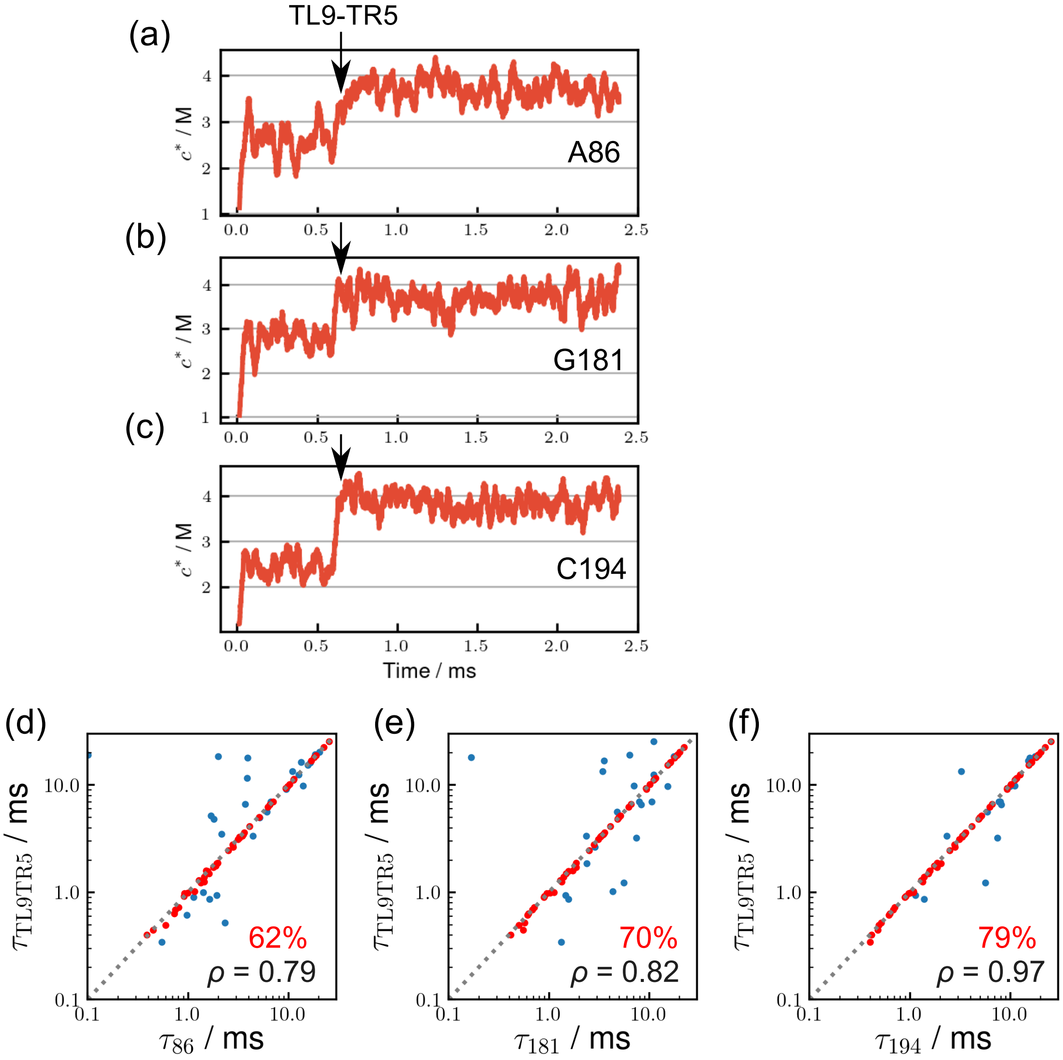


FIG. S20. **Coordinated formation of TL9-TR5 and $Mg^{2+}$ binding to A86, G181, and C194. (a-c)** $Mg^{2+}$ binding to (a) A86, (b) G181, and (c) C194 taken from the same simulation trajectory as Figure 3. The black arrows show the folding time of TL9-TR5. **(d-f)** Scatter plots of the first passage times of $Mg^{2+}$ binding to (d) A86, (e) G181, and (f) C194 versus the first passage time of the TL9-TR5 formation. In the majority of these trajectories, there is a high degree of specific coordinated $Mg^{2+}$ binding and TL9-TR5 formation.

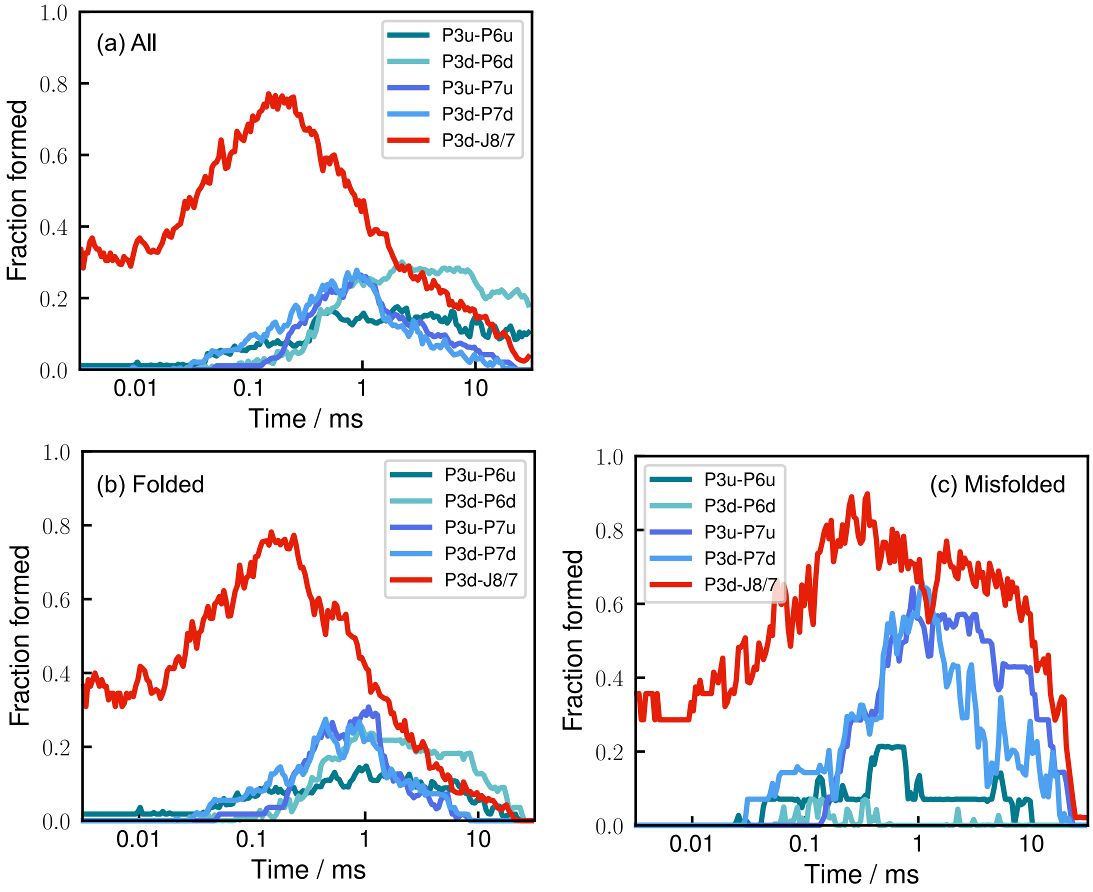


FIG. S21. **Formation of the five mispaired helices, which slows the folding of the ribozyme. (a)** Time-dependent fractions of mispaired helices averaged over all the trajectories. **(b, c)** Same as (a) except those correspond to trajectories that reach the folded state (b), and the misfolded (J8/7) state (c). See Figure S8 for the notation of the mispaired helices.
